# Epigenetic Reactivation of Lineage Differentiation to Target Leukemia

**DOI:** 10.64898/2026.07.08.737260

**Authors:** Shandon M. Amos, Chi-Chao Chen, Yichen Xiang, Keisuke Motoyama, Tania J. González-Robles, Varun Narendra, Grace Johnson, Han Teo Lee, Yu-Jui Ho, Ipek Celikoyar, Ziyang Ye, Shenghao Guo, Claire Glickman, Natalie O’Hearn, Oindrila Sarkar, Arianna Arroyo-Ortega, Tessa Devine, Michele Pagano, Kelly V. Ruggles, Francisco J. Sánchez-Rivera, Angela N. Koehler, Scott W. Lowe, Yadira M. Soto-Feliciano

## Abstract

Chromatin regulation critically influences gene expression and cancer progression, yet the functions of chromatin adaptors remain incompletely defined. Using focused CRISPR screening, we identified TRIM28, a multi-domain chromatin adaptor, as a dependency in acute leukemia, where its depletion impaired leukemia cell proliferation *in vitro* and *in vivo*, while activating neutrophil differentiation programs. Integrative transcriptomic and chromatin profiling revealed that TRIM28 acts as a co-repressor of neutrophil-associated loci independently of H3K9 methylation, and that TRIM28 loss drives terminal differentiation of leukemia cells into functionally mature neutrophil-like cells with reduced leukemic potential. We developed a selective small-molecule TRIM28 inhibitor that binds the TRIM28 PHD-bromodomain, phenocopies *TRIM28* loss across biochemical and cellular assays, exhibits low micromolar anti-leukemia activity, induces neutrophil differentiation, and synergizes with Menin inhibition. Together, these findings, spanning target discovery, mechanism of action, and chemical probe development, establish TRIM28 as a regulator of myeloid cell fate and a promising pro-differentiation therapeutic target in acute leukemia.

## Introduction

Chromatin and epigenetic dysregulation are fundamental drivers of tumorigenesis and cancer cell evolution^1,2^. Advances in this field have led to the development of therapies that target the molecular machinery controlling the epigenome^3^, yet clinical resistance remains a major challenge^4–7^. In addition to resistance mechanisms specific to individual targets, many cancers display phenotypic and lineage plasticity, enabling rapid transition into states that are independent of their original chromatin and transcriptional configurations^8^. These features highlight the need to uncover additional mechanisms of epigenetic dysregulation to enable more effective interventions. While most research has focused on enzymes that modify chromatin, growing evidence suggests that non-enzymatic structural components of chromatin complexes, such as scaffolds and adaptor proteins, represent a largely underexplored but promising class of Amos & Chen et al. 2026 (preprint) therapeutic targets, as illustrated by the clinical success of Menin-MLL inhibitors^9–11^. Identifying such components in cancer could open new avenues for therapeutic intervention.

Hematopoietic stem and progenitor cell-derived cancers, including leukemias, are particularly reliant on abnormal chromatin regulation for both initiation and maintenance of malignancy^12^. Differentiation therapies have achieved clinical success, notably in PML-RARα-driven acute promyelocytic leukemia, but remain limited to a small subset of genetically defined cases^13^. Recent work showed that simultaneous inhibition of epigenetic (LSD1) and signaling (WNT/GSK3) regulators can rewire lineage and interferon programs^14^. This highlights the many mechanistically distinct chromatin and transcriptional pathways that remain to be discovered and therapeutically exploited in leukemia. Discovering new mechanisms that restore leukemic cell differentiation and cell-cycle exit could significantly broaden the impact of epigenetic therapies.

Here, we identify TRIM28/KAP-1 as a previously unrecognized genetic dependency in acute leukemia. TRIM28 acts as a context-dependent chromatin adaptor15 by scaffolding leukemia-specific protein complexes that repress neutrophil differentiation programs. Chronic depletion of TRIM28 induces terminal myeloid differentiation of leukemia cells into quasi-functional neutrophil-like cells and reduces their proliferation. Building on these insights, we used a biochemical and cellular screening pipeline to discover and characterize a first-in-class small-molecule TRIM28 inhibitor that phenocopies key features of genetic *TRIM28* loss and selectively impairs leukemia cell fitness. These findings demonstrate that TRIM28 is required for leukemia cell proliferation and establish direct pharmacologic inhibition of TRIM28 as a pro-differentiation therapeutic strategy in acute leukemia.

## Results

### Chromatin-focused CRISPR screen identified TRIM28 as a leukemia-specific dependency

Chromatin and epigenetic factors drive leukemia development and maintenance^12,16^. To identify novel epigenetic regulators essential for leukemia cell growth, we performed a focused CRISPR loss-of-function screen using a mouse model of *MLL*-rearranged (*MLL*-r) leukemia, driven by the MLL-AF9 oncofusion^17^ **(Figure 1A)**. A mouse leukemia cell line expressing *Cas9* was transduced with a lentiviral single-guide RNA (sgRNA) library targeting ∼650 genes encoding chromatin regulatory proteins^4^. After approximately 12 population doublings, sgRNA representation in the cell population was assessed by high-throughput sequencing. We identified sgRNAs targeting ∼80 genes that were significantly depleted at the final time point (T_f_) from the cell population compared to the initial input (T_0_) **(Figure 1B)**. In addition to novel leukemia dependencies, several hits from the screen correspond to known leukemia dependencies, including *Setdb1*^18^, *Men1*^19–22^, inhibitors of which are now FDA-approved and marketed for acute leukemia; and *Brd4*^23,24^, inhibitors of which are currently under clinical investigation^25^.

**Figure 1.**
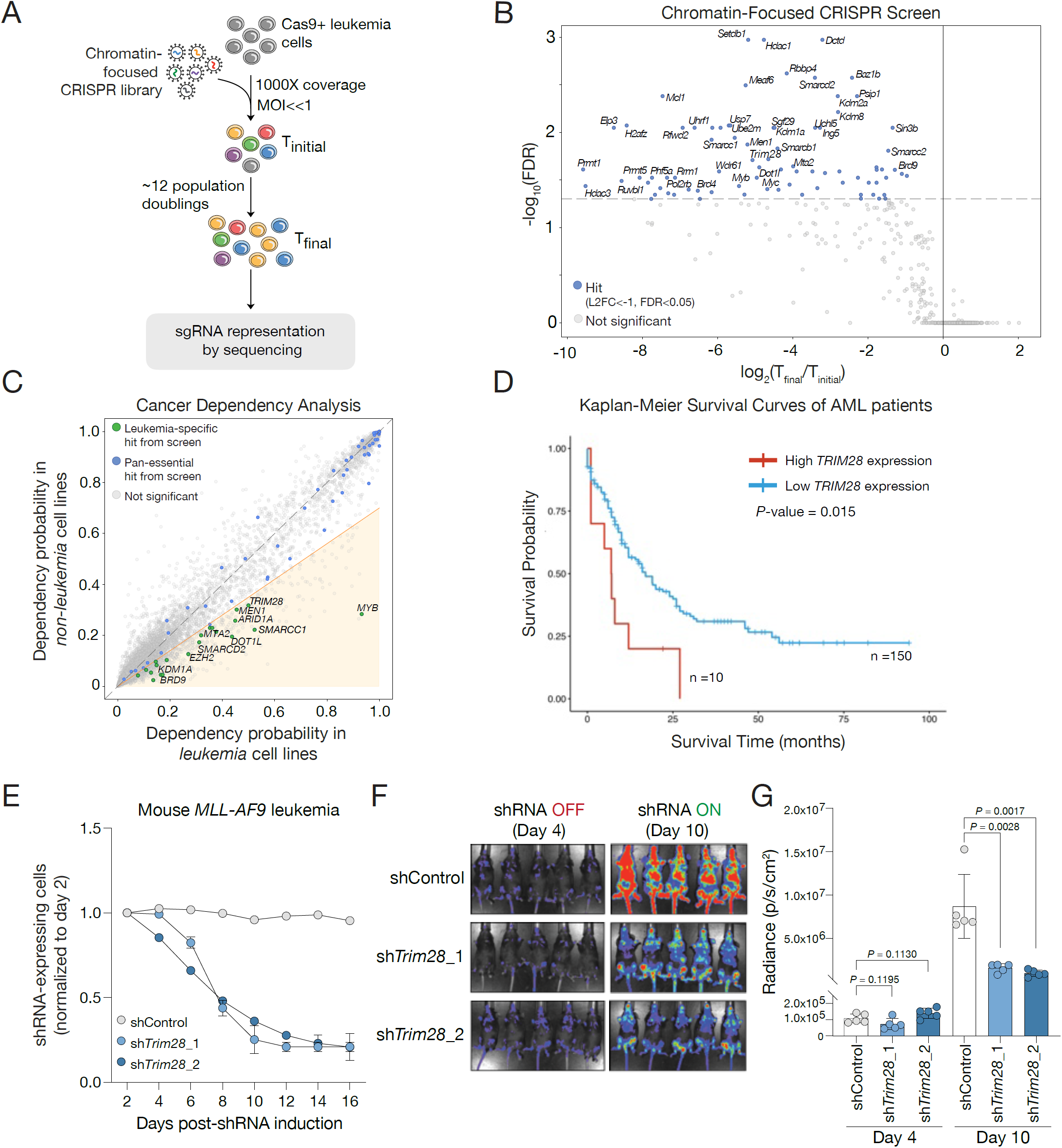
Chromatin-focused CRISPR screen identifies TRIM28 as a leukemia-specific dependency. (A) Experimental layout of the chromatin-focused CRISPR screen in mouse MLL-AF9 leukemia cells. (B) Volcano plot displaying log2(fold-change) for each gene tested in the chromatinfocused sgRNA library across screen replicates (n = 3). Significantly depleted genes, defined as log2(fold-change) < -1 and FDR < 0.05, are shown in blue. (C) Cancer dependency plot using the Broad Institute DepMap database, where the x-axis represents the dependency probability in leukemia cell lines (n = 118) and the y-axis represents the dependency probability in non-leukemia cell lines (n = 6,197). Hits from the chromatin-focused CRISPR screen (n = 81) are highlighted in blue if pan-essential and in green if leukemia-specific dependencies, defined as genes with a dependency probability ≥1.5-fold higher in leukemia cell lines than in all other cancer types, as indicated by the shaded yellow region. (D) Kaplan-Meier survival curves from TCGA AML patients stratified by high (n = 10) compared to low (n = 150) TRIM28 expression; P-value calculated by log-rank test. (E) Growth competition experiment in mouse MLL-AF9 leukemia cells engineered with doxycycline-inducible shRNA (non-targeting: shControl, Trim28-targeting: shTrim28_1 and shTrim28_2) co-expressing GFP. Cells were treated with 1 μg/mL doxycycline, and the proportion of GFP+ cells was quantified every 2 days over a 16-day period by flow cytometry (n = 3 replicates). (F) Syngeneic transplantation of mouse MLL-AF9 leukemia cells expressing Trim28 shRNAs followed by in vivo bioluminescence imaging (IVIS) of mice prior to shRNA induction (day 4 post-transplant) and after 1 week of knockdown (day 10). (G) Quantification of bioluminescence signal for shControl and shTrim28 groups, showing individual replicates (n = 5) and corresponding *p-*value

To identify leukemia-specific targets, we cross-referenced the genetic screen hits with data from the Broad Institute Cancer Dependency Map (DepMap)^26^, categorizing genes as either (1) pan-essential or (2) non-pan-essential across 2,000 human cancer cell lines. We defined leukemia-specific dependencies as genes with a dependency score at least 1.5 times higher in leukemia cell lines compared to all other cancer types, as indicated by the shaded yellow region **(Figure 1C)**. *TRIM28* emerged as a highly selective and significant dependency in acute leukemia of both myeloid and lymphoid lineages, clearly positioned off the diagonal in DepMap data and thus distinct from pan-essential genes **(Figure 1C)**. Analysis of acute myeloid leukemia (AML) patient survival data from The Cancer Genome Atlas (TCGA)^27^ further revealed that elevated *TRIM28* mRNA expression is significantly associated with worse patient outcomes, with distinct median survival differences between high- and low-expression groups **(Figure 1D)**. Importantly, the same analysis performed using *MEN1* expression (the most recently approved drug target in AML)^10,11^ revealed a similar but less significant survival trend **(Supplementary Figure 1A)**. Taken together, these data position TRIM28 as a promising therapeutic target in leukemia.

### Genetic knockdown of *TRIM28* inhibits leukemia proliferation

To investigate the functional consequences of *Trim28* loss in leukemia, we introduced a doxycycline-inducible *Trim28* knockdown system^28^ into the same mouse model of *MLL-r* leukemia^17^ used in our CRISPR screen **(Supplementary Figure 1B-C)**. We observed significant depletion of *Trim28*knockdown leukemia cells over time, measured by a reduction in GFP-positive (GFP^+^) cells **(Figure 1E, Supplementary Figure 1D)**. We expanded this analysis across multiple mouse models of leukemia, each driven by distinct genetic alterations, and observed significant leukemia cell depletion following *Trim28* knockdown **(Supplementary Figure 1E-H)**. Non-malignant mouse embryonic fibroblasts (MEFs) expressing identical constructs showed no proliferative defects after *Trim28* loss, confirming the leukemia-specific nature of this genetic dependency **(Supplementary Figure 1I)**.

To test whether this sensitivity is conserved in human cells, we knocked out *TRIM28* in several leukemia cell lines. Regardless of driver mutation, sg*TRIM28*-expressing cells were robustly depleted from the population **(Supplementary Figure 2A-F)**. We next generated a doxycycline-inducible *TRIM28* knockdown system in human AML cells (MOLM13) **(Supplementary Figure 3A)**, and as in mouse cell lines, the proportion of sh*TRIM28*- expressing cells significantly decreased over time **(Supplementary Figure 3B)**. Expression of full-length mouse *Trim28* cDNA, which is resistant to the human-specific sh*TRIM28* **(Supplementary Figure 3C)**, fully rescued the proliferation defect in these leukemia cells **(Supplementary Figure 3D)**.

To validate the dependency of leukemia cells on *Trim28 in vivo*, we transplanted mouse AML cells into immunocompetent syngeneic recipient mice **(Supplementary Figure 4A)**. *Trim28* knockdown after engraftment significantly impaired leukemia progression, as evidenced by reduced bioluminescent signal intensity and fewer circulating leukemia cells in peripheral blood **(Figure 1F-G, Supplementary Figure 4B)**. This was accompanied by significantly lowered white blood cell counts and prolonged survival of recipient mice **(Supplementary Figure 4C-D)**. Using human leukemia cells transplanted into immunodeficient mice, we also observed significantly lower leukemia burden in mice and extended survival following *TRIM28* knockdown **(Supplementary Figure 4E-G)**. Together, these findings demonstrate that AML cells are highly dependent on TRIM28 both *in vitro* and *in vivo*. This previously unrecognized vulnerability is conserved across mouse and human leukemia models and is independent of specific oncogenic drivers (e.g. *MLL-r*).

### TRIM28 PHD-bromodomain is critical for its function in leukemia

TRIM28 acts as a chromatin scaffold, assembling protein complexes that regulate gene expression and heterochromatin formation through coordinated activity of its N-terminal RBCC (TRIM motif)^29–31^ and C-terminal epigenetic effector domains^32,33^. It is essential for embryonic development, with homozygous knockout resulting in early post-implantation lethality^34^, while loss in adult tissues has minimal to no effect^35^. The fact that leukemia depends on a non-catalytic chromatin scaffold, rather than a single enzymatic subunit reveals a mechanistically distinct mode of epigenetic control and highlights scaffold proteins as an attractive class of targets for therapeutic intervention.

To identify the specific TRIM28 domains required for leukemia maintenance, we performed a CRISPR-based saturation mutagenesis tiling screen **(Figure 2A)**. This strategy relies on the established principle that insertion-deletion (indel) mutations outside essential, well-structured domains are generally tolerated, whereas even in-frame alterations are often deleterious within conserved, folded domains^36^. A pooled tiling sgRNA library spanning the entire *TRIM28* coding region was introduced into a human leukemia cell line (MOLM13). After ∼20 days of culture, sgRNA representation was deconvoluted by high-throughput sequencing at the initial (T_0_) and final (T_f_) time points. Significant dependency regions were identified using the CKHS saturation mutagenesis depletion score, calculated from the Z-score of each sgRNA^37^. We observed significant depletion of sgRNAs targeting the coiled-coil, HP1-binding domain, and bromodomain **(Figure 2B)**. The centrally located coiled-coil domain of TRIM28, spanning approximately 400 amino acids, is thought to be responsible for mediating the protein’s homodimerization and stability^38^. This function is supported by the predicted AlphaFold 3 model^39,40^ **(Figure 2C)**. The HP1-binding motif consists of a short stretch of residues within the intrinsically disordered region (IDR), which also harbors the predicted nuclear localization sequence41. A strong depletion of sgRNAs targeting the approximately 80 amino acid bromodomain (BD) at the C-terminus was observed. This suggests that this previously uncharacterized region may play a critical role in TRIM28 function in leukemia maintenance **(Figure 2B-C)**.

**Figure 2.**
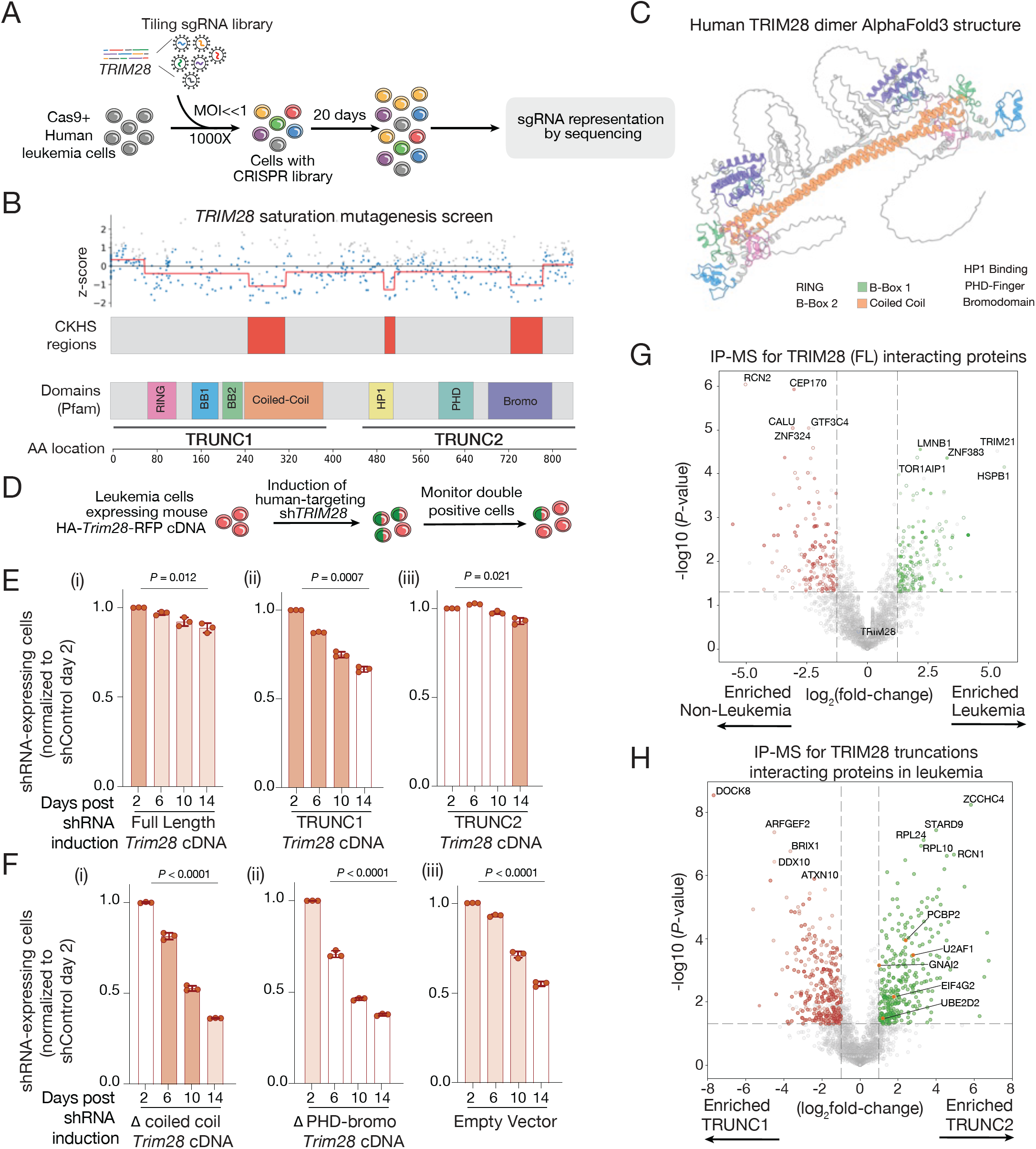
TRIM28 PHD-bromodomain is critical for its function in leukemia. **(A)** Schematic of the CRISPR-mediated *TRIM28* saturation mutagenesis screen in MOLM13 leukemia cells. **(B)** (Top) sgRNA depletion scores across the *TRIM28* coding sequence; all sgRNAs shown in grey, significant guides (*FDR* < 0.05) in blue, and rolling average in red. (Middle) CRISPR knockout hyper-sensitive (CKHS) region scores. (Bottom) TRIM28 Pfam domain annotation. **(C)** AlphaFold 3 structure of human TRIM28 homodimer with domains highlighted. **(D)** Schematic of doxycycline-inducible shRNA-mediated competition rescue experiments. **(E)** Competition rescue experiment in MOLM13 cells constitutively expressing HA-tagged mouse *Trim28* cDNA (resistant to human-targeting shRNAs), engineered with doxycycline-inducible shRNAs (non-targeting shControl; *TRIM28*-targeting sh*TRIM28*_1 and sh*TRIM28*_2) co-expressing *GFP*. GFP^+^ cell proportion was quantified every 4 days over 14 days by flow cytometry. Panels show rescue with (i) full-length (FL), (ii) N-terminus-leucine zipper (TRUNC1), and (iii) HP1-binding domain-C-terminus (TRUNC2) mouse *Trim28* cDNA. Significance was assessed by comparing GFP^+^ proportions at day 2 vs. day 14. **(F)** As in panel E, with panels showing rescue with mouse *Trim28* cDNA containing (i) an internal leucine zipper deletion, (ii) a PHD-bromodomain deletion, or (iii) empty vector. **(G)** Volcano plot of TRIM28-interacting proteins in MOLM13 leukemia vs. HEK293T non-leukemia cells by co-IP/MS (n = 4 replicates). Proteins enriched in leukemia cells (log_2_(fold-change) > 1, *P*-value < 0.05; 199 proteins) are shown in green; those enriched in non-leukemia cells (log_2_(fold-change) < −1, *P*-value < 0.05; 188 proteins) in red. **(H)** Volcano plot of TRIM28-interacting proteins in TRIM28-TRUNC2-vs. TRIM28-TRUNC1-expressing MOLM13 cells by co-IP/MS (n = 4 replicates). Proteins enriched in TRUNC2-expressing cells (log_2_(fold-change) > 1, *P*-value < 0.05; 310 proteins) are shown in green; those enriched in TRUNC1-expressing cells (log_2_(fold-change) < −1, *P*-value < 0.05; 317 proteins) in red.

We performed complementary structure-function analyses to define the domains necessary and sufficient for rescuing proliferative capacity following *TRIM28* knockdown in leukemia. One series of deletion mutants divided the protein into two segments: TRUNC1, encompassing the N-terminus through the coiled-coil domain, and TRUNC2, spanning the HP1-binding motif through the C-terminus **(Figure 2B)**. A second set of variants carried internal deletions within either the coiled-coil domain or the PHD-bromodomain, referred to as Δcoiled-coil and ΔPHD-bromo, respectively. Human leukemia cells were engineered to express mouse *Trim28* cDNA encoding these variants, each fused to an N-terminal HA tag and co-expressed with RFP to enable monitoring via flow cytometry **(Figure 2D, Supplementary Figure 5A)**. These truncations showed localization to the nuclear and cytoplasmic cellular compartments as was also observed with the full-length protein **(Supplementary Figure 5B)**.

Upon *TRIM28* knockdown, we found that TRUNC2 fully rescued the proliferation defect, similar to full-length TRIM28 **(Figure 2E)**. In contrast, TRUNC1, Δcoiled-coil, and ΔPHD-bromo deletion constructs failed to restore proliferation **(Figure 2E-F)**. We hypothesized that the inability of the Δcoiled-coil construct to rescue function results from steric interference between the PHD-bromodomain and the RING and B-Box1/2 (BB1/2) domains, producing a shorter, potentially more rigid, and dysfunctional protein. Structural modeling of these TRIM28 variants using AlphaFold 3^40^ supports this hypothesis, where the Δcoiled-coil variant yielded the lowest predicted TM-score (pTM = 0.35), consistent with substantial structural disruption, whereas TRUNC1 (pTM = 0.55), TRUNC2 (pTM = 0.45), and ΔPHD-bromo (pTM = 0.43) all showed higher predicted fold confidence **(Supplementary Figure 6A-D)**.

### TRIM28 assembles leukemia-specific complexes associated with cell identity

TRIM28 is a chromatin scaffold protein; we then asked whether it assembles unique protein complexes in leukemia. We performed co-immunoprecipitation followed by mass spectrometry (co-IP/MS) in human leukemia and non-leukemia cells expressing *TRUNC1, TRUNC2*, or full-length *Trim28*. We defined high-confidence interactors by combining data from IPs of both endogenous TRIM28 and exogenous HA-tagged full-length TRIM28 in each cell type before comparing interactions across contexts. We identified 199 high-confidence TRIM28 interactors in leukemia and 188 in non-leukemia cells **(Figure 2G-H)**. Enrichment of proteins in the TRIM28 immunoprecipitation did not correlate with the expression levels of the corresponding genes (Pearson’s *r* = -0.126), indicating that differences in protein interactions are not simply driven by mRNA abundance in each cell type **(Supplementary Figure 7A)**. Pathway analysis of TRIM28-interacting proteins revealed distinct biological processes associated with leukemia versus non-leukemia cells **(Supplementary Figure 7B)**. In leukemia cells, TRIM28-interacting proteins were enriched for pathways related to signaling and immune cell regulation, including NF-κB signaling, cell surface receptor-mediated signaling, and tumor necrosis factor **(Supplementary Figure 7B)**. In contrast, TRIM28-associated proteins in non-leukemia cells were enriched for processes related to ribosome biogenesis, RNA metabolism, and translation. These results indicate that TRIM28 assembles functionally distinct protein complexes depending on cellular context, shifting from roles in RNA biogenesis in non-leukemic cells to engagement with signaling and immune pathways in leukemia.

TRIM28-TRUNC2 is necessary and sufficient to complement *TRIM28* knockdown, whereas TRIM28-TRUNC1 only partially rescues the phenotype. To identify separation-of-function complexes within TRIM28, we analyzed the interactome of each segment individually. Surprisingly, *TRUNC1*-expressing cells recovered TRIM28-containing protein complexes linked to processes resembling those of full-length TRIM28 in leukemia, including leukocyte migration and myeloid cell differentiation **(Supplementary Figure 7C)**. In contrast,

*TRUNC2*-expressing leukemia cells were enriched for proteins involved in pathways typical of non-leukemic contexts, such as mRNA processing, splicing, and ribosome biogenesis **(Supplementary Figure 7C)**. Together with the complementation data, these results suggest that the regulatory functions contained within TRUNC2 are more essential and act downstream in leukemia cells compared to TRUNC1. TRUNC2 supports scaffolding and recruitment of epigenetic silencing complexes such as SETDB1^32^ and SIN3A^42^, while TRUNC1 exhibits E3 ubiquitin ligase activity^43^ that targets client proteins for degradation **(Supplementary Figure 7D)**. Collectively, these findings suggest that TRIM28 may maintain leukemia cell identity by both degrading immune effector proteins and repressing transcriptional regulators of myeloid differentiation. To investigate co-dependency among TRIM28 interacting partners, we integrated our proteomics data with genetic dependency profiles from the Cancer Dependency Map (DepMap)^26^. Among the strongest correlated genes were *MED23, CSDE1, STAG2*, and *ZNF148* **(Supplementary Figure 8A)**. Network analysis of TRIM28 co-dependencies revealed a dense interaction hub centered on TRIM28, with many top co-dependent genes involved in RNA processing (*CSDE1, DHX29*), transcriptional regulation (*MED23, SP1, ARX*), and chromatin organization (*STAG2, SMARCD1*) **(Supplementary Figure 8B)**. A subset of these genes overlapped with known physical interactors annotated in BioGRID^44^, such as TIPARP, ALDH9A1, and DHX29, reinforcing the close alignment between TRIM28’s biochemical and genetic interaction networks. To link these genetic dependencies to TRIM28’s biochemical functions, we compared TRIM28 TRUNC2-associated proteins identified in leukemia cells with genes positively co-dependent with *TRIM28* in DepMap. This analysis revealed a substantial overlap, with 1,071 shared factors (56% of all TRUNC2 interactors) out of 1,906 unique TRUNC2 interactors and 1,863 unique TRIM28 myeloid co-dependencies. This highlights a large group of proteins that both physically associate with TRIM28’s C-terminal domains in leukemia and display coordinated essentiality with *TRIM28* across myeloid neoplasm cell lines **(Supplementary Figure 8C)**.

Collectively, these data suggest that TRIM28 promotes leukemia proliferation via context-specific interactions mediated predominantly by its C-terminal PHD-bromodomain.

### TRIM28 suppresses neutrophil-associated gene programs in leukemia

To define TRIM28-regulated gene programs in leukemia, we transcriptionally profiled cells after *Trim28* knockdown. We generated single-cell clones of mouse MLL-AF9 leukemia cells expressing two doxycycline-inducible shRNAs targeting *Trim28* and confirmed on-target depletion **(Supplementary Figure 9A-B)**. *Trim28* knockdown resulted in widespread transcriptional changes, predominantly upregulation **(Figure 3A, Supplementary Figure 9C-D)**. Differential expression analysis identified 2,470 genes significantly altered upon *Trim28* knockdown, with 1,716 upregulated and 756 downregulated **(Figure 3A)**. *K*-means clustering of the RNA-Seq data identified five gene clusters defined by the magnitude and direction of TRIM28-dependent expression changes **(Figure 3B)**. Cluster 1 (271 genes) contained rapidly upregulated genes enriched for positive regulation of transcription from RNA Polymerase II promoter. Cluster 2 was the largest and most rapidly responding cluster containing 706 TRIM28-repressed genes, which were strongly enriched for neutrophil degranulation pathways by gene ontology (GO) analysis^45^ **(Figure 3B)**. The early induction of transcriptional regulators (Cluster 1) followed by the upregulation of neutrophil-associated effector genes (Cluster 2) is consistent with a hierarchical activation model in which TRIM28 loss first de-represses key transcription factors that subsequently drive a neutrophil differentiation program.

**Figure 3.**
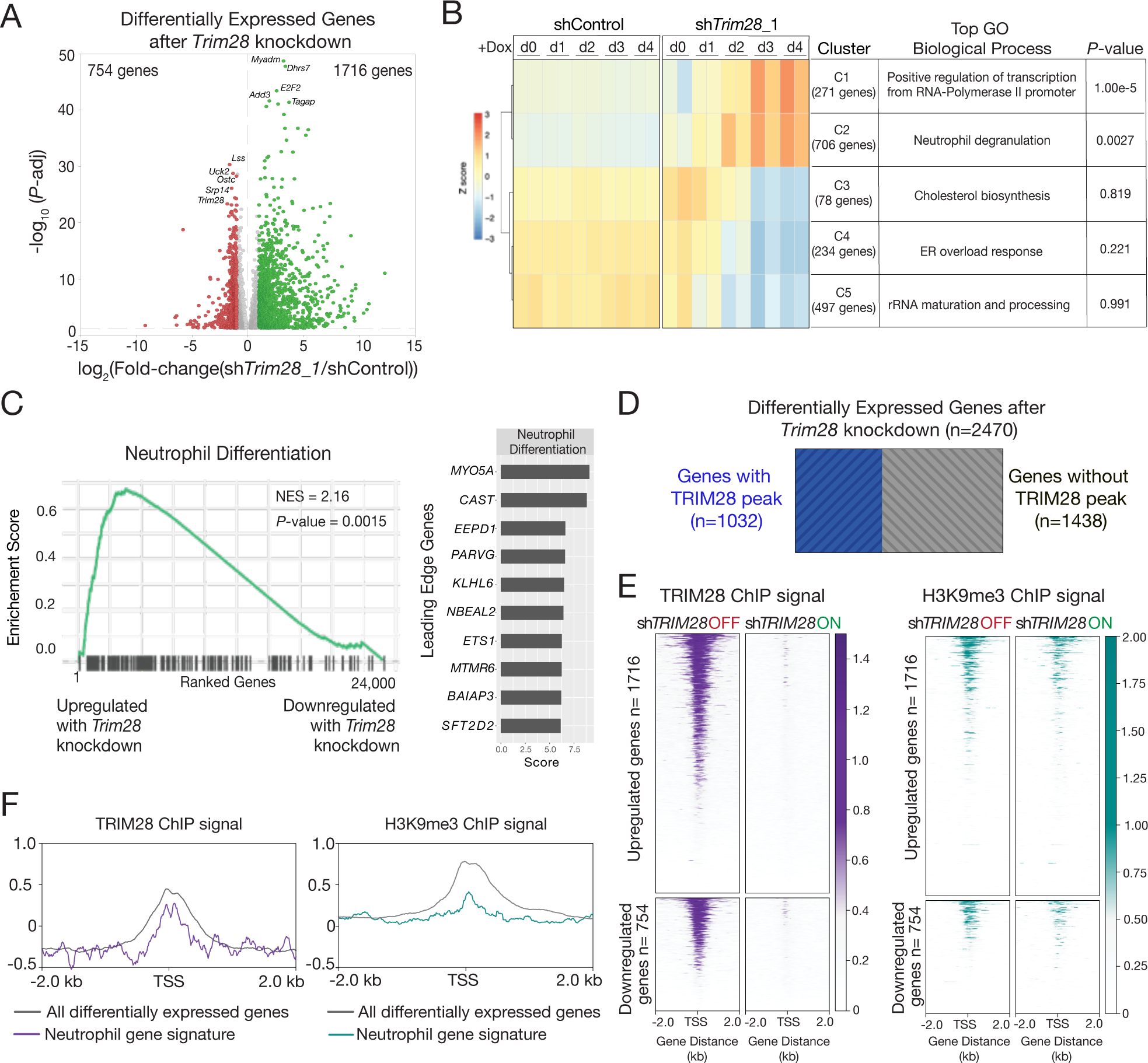
TRIM28 loss results in transcriptional induction of neutrophil-associated gene programs. **(A)** Volcano plot of differentially expressed genes measured by RNA-Seq in mouse leukemia cells after 4 days of *Trim28* knockdown (sh*Trim28*_1) compared to a non-targeting control (n = 3 replicates). Genes significantly upregulated following *Trim28* knockdown (log_2_(fold-change) > 1 and -log_10_(*P*-value) > 1.3, n = 1,716 genes) are shown in green, and genes significantly downregulated (log_2_(fold-change) < -1 and -log_10_(*P*-value) > 1.3, n = 754 genes) are shown in red. **(B)** (Left) Heatmap of expression for differentially expressed genes (*Z*-score; |log_2_(fold-change)| > 1 and *P*-value < 0.05) before and 1, 2, 3, and 4 days after *Trim28* knockdown (sh*Trim28*_1). *K*-means clustering grouped genes into five clusters and the top-enriched Gene Ontology (GO) biological process term for each cluster with its associated *P*-value are shown on the right. **(C)** GSEA plot showing enrichment of a neutrophil differentiation gene signature (GSE27786) following 4 days of *Trim28* knockdown (sh*Trim28*_1) in mouse leukemia cells. The normalized enrichment score (NES) and *P*-value are indicated, and the top 10 leading-edge genes are shown in the bar plot to the right. **(D)** Bar plot showing the proportion of differentially expressed genes (RNA-Seq) that overlap or do not overlap with TRIM28 chromatin occupancy (ChIP-Seq). **(E)** (Left) Heatmap of TRIM28 ChIP-Seq signal at genes upregulated (log_2_(fold-change) > 1 and -log_10_(*P*-value) > 1.3, n = 1,716 genes) and downregulated (log_2_(fold-change) < -1 and -log_10_(*P*-value) > 1.3, n = 754 genes) after *Trim28* knockdown (sh*Trim28*_1). (Right) Heatmap of H3K9me3 ChIP-Seq signal at the same genes. **(F)** (Left) Metagene plot of TRIM28 chromatin occupancy at all differentially expressed genes (grey) and at genes within the neutrophil differentiation gene set (purple). (Right) Metagene plot of H3K9me3 chromatin occupancy showing all differentially expressed genes (grey) and neutrophil differentiation signature genes (teal).

Gene set enrichment analysis (GSEA)^46^ confirmed that the neutrophil differentiation gene signature was the most significantly enriched program among genes upregulated upon *Trim28* knockdown **(Figure 3C)**. Broader pathway analysis showed significant enrichment of pathways related to neutrophil differentiation, neutrophil degranulation, myeloid cell development, and the CEBPα network, as well as interleukin signaling pathways **(Figure 3C, Supplementary Figure 9E)**. A complementary analysis combining both hairpins further confirmed this observation **(Supplementary Figure 9F)**. These data suggest that TRIM28 acts as a critical repressor of neutrophil differentiation-associated gene programs in leukemia.

To examine if these effects were conserved in humans, we performed transcriptional profiling of the human leukemia cell line MOLM13 following *TRIM28* knockdown **(Supplementary Figure 10A-C)**. We observed robust transcriptional activation of neutrophil differentiation pathways **(Supplementary Figure 10D-E)**, as in mouse leukemia cells. Because TRIM28 is known to repress endogenous retroelements^47,48^, we examined poly-adenylated transposable elements (TE)^49^ in *TRIM28*-depleted leukemia cells to determine whether TE activation underlies the observed transcriptional changes. We observed minimal changes, with only one element (*LTR4:ERV1:LTR*) showing moderate upregulation after 4 days of *TRIM28* knockdown **(Supplementary Figure 10F)**. Together, these findings suggest that TRIM28 maintains the leukemic state by repressing neutrophil differentiation programs, independent of its role in TE silencing.

### TRIM28 employs a non-canonical mechanism to repress neutrophil differentiation in leukemia

Our transcriptional analysis revealed a previously unappreciated role for TRIM28 as a negative regulator of neutrophil differentiation gene programs in leukemia. Canonically, TRIM28 mediates transcriptional repression through SETDB1, which deposits H3K9me3 to promote heterochromatin formation^33,50^. To test whether this pathway underlies TRIM28-dependent suppression of neutrophil differentiation in leukemia, we performed ChIP-Seq for TRIM28 and H3K9me3 in mouse MLL-AF9 leukemia cells. We identified over 13,000 high-confidence TRIM28 ChIP peaks that were lost or markedly reduced after Trim28 knockdown **(Supplementary Figure 11A)**, of which 1,032 mapped to genes differentially expressed upon knockdown **(Figure 3D)**. TRIM28 ChIP peaks were present at both upregulated and downregulated genes, indicating that TRIM28 can function as either a co-repressor or co-activator depending on chromatin context and interacting partners **(Figure 3E)**. However, TRIM28 occupancy and H3K9me3 enrichment were not uniformly correlated at these loci **(Supplementary Figure 11B-C)**. At upregulated genes, TRIM28 ChIP signal was robustly enriched at TSSs and significantly reduced upon knockdown, yet H3K9me3 showed a comparatively modest decrease **(Supplementary Figure 11B)**; a similar pattern held at downregulated genes **(Supplementary Figure 11C)**. This dissociation between TRIM28 binding and H3K9me3 deposition suggested that TRIM28 represses a subset of its target genes through mechanisms independent of its canonical SETDB1-H3K9me3 axis.

To distinguish SETDB1-dependent from SETDB1-independent repression, we intersected upregulated genes with the presence or absence of overlapping H3K9me3 peaks **(Supplementary Figure 11D)**. 396 upregulated genes displayed decreased H3K9me3 signal after TRIM28 loss, consistent with SETDB1-mediated repression, whereas 337 upregulated genes lacked detectable H3K9me3 peaks, indicating SETDB1-independent repression **(Supplementary Figure 11D)**. Gene ontology analysis revealed enrichment for GPCR signaling pathways in both classes, but only the SETDB1-independent repression group was enriched for neutrophil degranulation and innate immune signatures **(Supplementary Figure 11E-F)**. This suggests that, although TRIM28 can repress transcription via canonical H3K9me3, its role in suppressing neutrophil differentiation in leukemia involves additional, SETDB1-independent mechanisms of chromatin regulation.

Analysis of TRIM28 and H3K9me3 average ChIP signals at neutrophil-associated gene signatures^51^ showed robust TRIM28 occupancy but minimal H3K9me3 enrichment at these loci **(Figure 3F)**. At the single-gene level, TRIM28 occupied the promoters of *Cebpa, Gfi1*, and *Spi1*, transcription factor loci critical for neutrophil lineage differentiation^52,53^, and this enrichment was markedly reduced upon knock-down, whereas H3K9me3 was either absent or minimal at the same loci **(Supplementary Figure 12A)**. Expression of *Cebpa, Gfi1*, and *Spi1* was significantly upregulated upon *Trim28* loss **(Supplementary Figure 12B)**. These data establish that TRIM28 directly binds the promoters of master neutrophil transcription factors in mouse leukemia cells and represses their expression independently of H3K9me3.

To determine whether this non-canonical mechanism is conserved in human leukemia, we performed CUT&RUN for TRIM28, H3K9me3, and the repressive histone modification H3K27me3 in human leukemia cells following *TRIM28* knockdown. At the *CEBPA, GFI1*, and *SPI1* promoters, TRIM28 was enriched and lost upon knockdown **(Supplementary Figure 12C)**. H3K9me3 was undetectable at all three loci; instead, H3K27me3 colocalized with TRIM28 and was lost upon knockdown **(Supplementary Figure 12C)**, implicating PRC2 rather than SETDB1 in TRIM28-mediated repression at these neutrophil-specifying lineage loci. Extending this analysis genome-wide, CUT&RUN quantification across all neutrophil differentiation gene promoters confirmed a robust reduction in TRIM28 signal, a significant decrease in H3K27me3, and no change in H3K9me3 upon *TRIM28* knockdown; notably, H3K4me3 increased at these loci, consistent with transcriptional de-repression/activation **(Supplementary Figure 12E)**. RNA-Seq confirmed concordant upregulation of *CEBPA/E, GFI1*, and *SPI1* **(Supplementary Figure 12D)**, and immunoblotting of nuclear extracts at day 6 following *TRIM28* knockdown verified that this de-repression extended to protein levels, with elevated CEBPA, GFI1, and SPI1 relative to day 0 and control cells **(Supplementary Figure 12F)**.

To more directly test whether TRIM28 cooperates with PRC2 at these loci, we performed co-immunoprecipitation using Alfa-tagged TRIM28 and confirmed a biochemical interaction between TRIM28 and EZH2 in leukemia cells **(Supplementary Figure 12G)**. Consistent with functional cooperation, EZH2 inhibition alone (tazemetostat) had little effect on SPI1 protein expression, but combining sh*TRIM28* with tazemetostat produced maximal SPI1 induction by immunoblot, while *TRIM28* knockdown alone (doxycycline) yielded an intermediate response **(Supplementary Figure 12H)**. These results demonstrate that TRIM28 functions as a co-repressor of neutrophil differentiation programs in leukemia independently of H3K9me3. Instead, TRIM28 may repress transcription in collaboration with the PRC2 complex, suggesting a novel role distinct from its well-characterized transcriptional repression of endogenous retroelements.

### TRIM28 loss leads to differentiation of leukemia into neutrophil-like cells

We found that TRIM28 sustains acute myeloid leukemia (AML) by transcriptionally repressing neutrophil differentiation genes **(Figure 3)**. Neutrophils, key effectors of innate immunity, constitute the most abundant and rapidly renewing cell population in the immune system^54,55^. Directed induction of AML cell differentiation into these short-lived neutrophils has been proposed to reduce relapse and prolong therapeutic responses, making neutrophil lineage commitment an attractive therapeutic endpoint^56^. To test whether TRIM28 depletion drives terminal neutrophil-like differentiation and thereby mediates leukemia cell elimination, we analyzed cell surface expression of CD16^57,58^ and CD66b^52,54^ by flow cytometry. After 12 days of *TRIM28* depletion, leukemia cells exhibited significantly increased CD66b expression, and a subset also expressed CD16, indicating progression toward a mature neutrophil-like state **(Figure 4A-B)**. Wright-Giemsa staining showed multilobed nuclei and granular cytoplasm typical of neutrophils^59^ **(Figure 4C)**. *TRIM28* knockdown did not increase apoptosis, indicating that terminal differentiation, not cell death, is the predominant cell fate **(Supplementary Figure 13A-B)**. Both the impaired proliferation and induction of neutrophil-like differentiation caused by *TRIM28* knockdown were fully rescued in human leukemia cells stably expressing a shRNA-resistant mouse *Trim28* cDNA **(Figure 4A-B, Supplementary Figure 3C-D)**.

**Figure 4.**
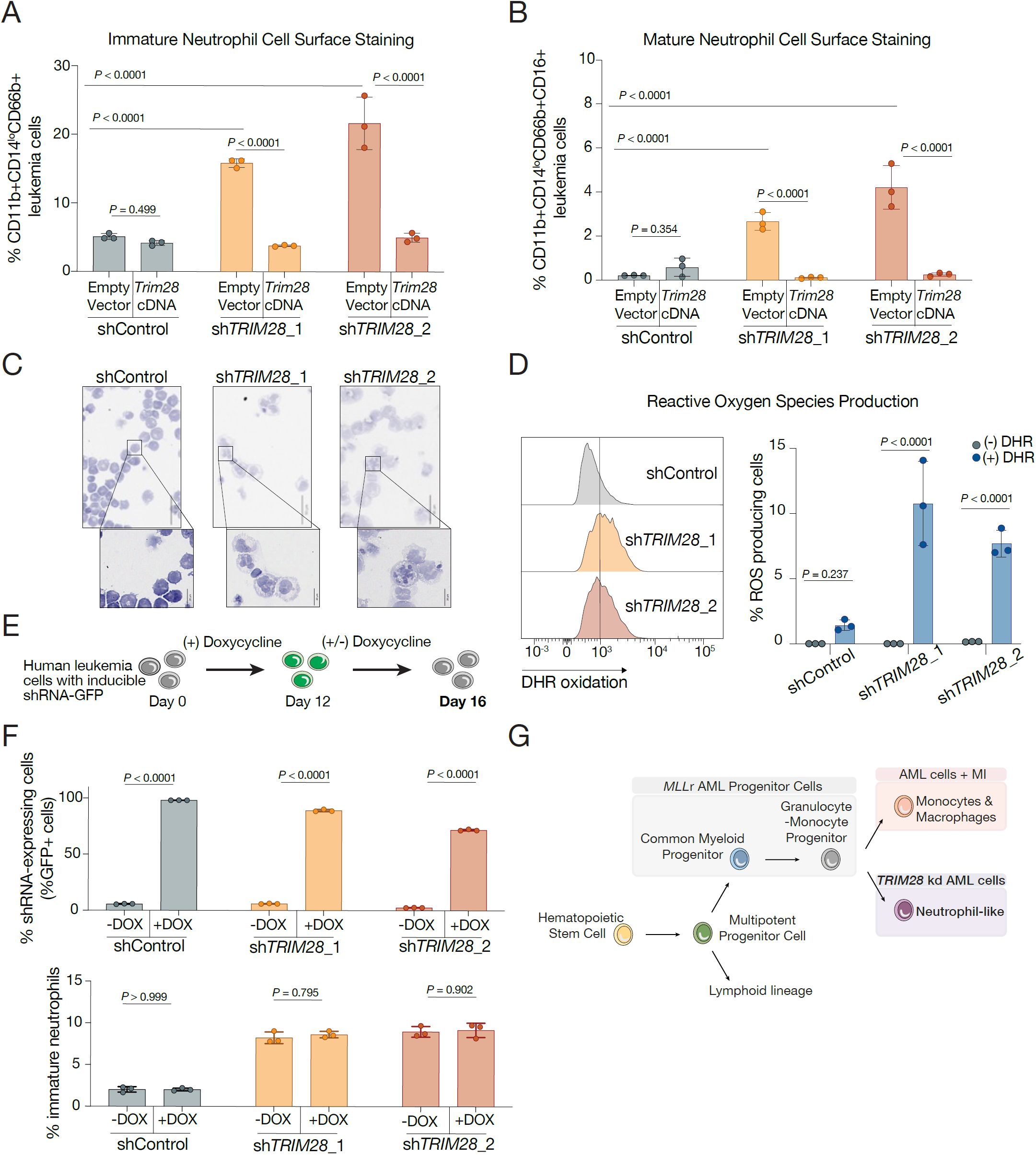
TRIM28 depletion induces irreversible neutrophil-like differentiation in leukemia cells. **(A)** Cell surface staining for immature neutrophils (CD11b^+^, CD14^lo^, CD66b^+^) in MOLM13 leukemia cells constitutively expressing HA-tagged full-length mouse *Trim28* cDNA, which is resistant to the human-targeting shRNAs, or empty vector control after 12 days of doxycycline-induced knockdown (n = 3 replicates). **(B)** Cell surface staining for mature neutrophils (CD11b^+^, CD14^lo^, CD66b^+^, CD16^+^) under the same conditions as in Figure 4A (n = 3 replicates). **(C)** Giemsa-Wright staining of MOLM13 leukemia cells after 12 days of doxycycline-induced shRNA expression. **(D)** (Left) Representative histogram of dihydrorhodamine 123 (DHR) oxidation in MOLM13 leukemia cells after 12 days of doxycycline-induced shRNA expression. (Right) Quantification of the percentage of DHR^+^ cells (n = 3 replicates). **(E)** (Top) Schematic of the experiment testing reversibility of *TRIM28* knockdown-induced phenotypes. MOLM13 cells with doxycycline-inducible shRNAs were treated with doxycycline for 12 days, then cultured with or without doxycycline for an additional 4 days before analysis at day 16. (Bottom) Proportion of GFP^+^ MOLM13 cells after removal or maintenance of doxycycline (shControl, sh*TRIM28*_1, sh*TRIM28*_2); n = 3 replicates. **(F)** Proportion of immature neutrophils (CD11b^+^, CD14^lo^, CD66b^+^) in MOLM13 cells after removal or maintenance of doxycycline under the same conditions as in Figure 4E (n = 3 replicates). **(G)** Schematic of the role of TRIM28 in regulating leukemia cell identity and neutrophil-like differentiation.

Normal activated neutrophils can generate reactive oxygen species (ROS) and form neutrophil extracellular traps (NETs), as part of the innate immune response^60^. To assess whether *TRIM28*-depleted leukemia cells acquired similar functional capabilities, cells were stimulated with granulocyte colony-stimulating factor (G-CSF) to mimic innate immune activation, and oxidative activity was measured using a dihydrorhodamine (DHR) oxidative burst assay^58,61^. Consistent with neutrophil-like functionality, only *TRIM28*-deficient cells displayed a significant oxidative burst **(Figure 4D)**. These cells also showed increased formation of NETs, evidenced by enhanced surface staining for myeloperoxidase (MPO) and histone H3^62^ **(Supplementary Figure 13C)**. These functional assays demonstrate that leukemia cells undergoing differentiation in response to *TRIM28* knockdown acquire functional hallmarks of mature neutrophils^63^.

Terminally differentiated neutrophils are short-lived cells, and their rapid clearance is predicted to be a desirable outcome during anti-leukemia therapy^56^. To determine whether the TRIM28 loss-driven neutrophil differentiation program is irreversible, doxycycline was withdrawn from cell cultures for 4 days following 12 days of TRIM28 knockdown **(Figure 4E)**. Although sh*TRIM28* expression declined, as shown by reduced GFP signal and restored endogenous TRIM28 protein levels **(Figure 4F, Supplementary Figure 13D-E)**, the proportion of neutrophil-like cells remained unchanged after doxycycline removal **(Figure 4F)**. These findings indicate that TRIM28 loss triggers a terminal and irreversible differentiation of leukemia cells into neutrophil-like cells **(Figure 4G)**.

Previous studies have shown that small-molecule inhibition of Menin-MLL induces monocyte and macrophage-associated transcriptional signatures in leukemia cells^22,64,65^. To evaluate the effects of simultaneous *TRIM28* knockdown and Menin-MLL inhibition, human leukemia cells were treated with a Menin inhibitor (MI-503) alongside *TRIM28* knockdown for 6 days **(Supplementary Figure 14A-B)**. Combined *TRIM28* knockdown and Menin inhibition significantly increased CD11b expression relative to Menin inhibition alone **(Supplementary Figure 14C-D)**. Cells receiving both perturbations exhibited surface markers characteristic of immature neutrophils rather than macrophages **(Supplementary Figure 14E-F)**. These results suggest that combined inhibition of TRIM28 and Menin exerts an additive effect in promoting myeloid differentiation, with a pronounced bias toward neutrophil-like maturation **(Figure 4G)**.

Together, these results show that TRIM28 loss drives leukemia cells into irreversible, functionally mature neutrophil-like differentiation, a therapeutically favorable outcome^56^. This work reveals TRIM28 as a critical regulator of myeloid cell fate and highlights co-targeting of TRIM28 and Menin as a promising strategy to more effectively promote differentiation and suppress leukemia cell proliferation.

### Novel TRIM28 inhibitor impairs leukemia maintenance and induces neutrophil differentiation

Our results identified TRIM28 as a selective dependency in leukemia that represses neutrophil differentiation programs. We next sought to pharmacologically target TRIM28 and define the consequences of its inhibition in leukemia cells.

We performed a high-throughput screen of 65,000 small molecules against the human TRIM28 PHD-bromodomain^66^, identifying primary binders by small-molecule microarray^67^ **(Supplementary Figure 15A)**. Iterative reproducibility and counter-screens followed by proliferation and differentiation assays narrowed the candidate set to a single lead compound, KI-T28-03 **(Figure 5A, Supplementary Figure 16A-B)**. In human leukemia cells, KI-T28-03 markedly reduced proliferation over 6 days, comparable to the Menin inhibitor MI-503, whereas most other candidates had little effect **(Figure 5B, Supplementary Figure 16B)**. In parallel, KI-T28-03 uniquely increased CD11b expression, indicating myeloid differentiation **(Figure 5C)**. Thermal-shift profiling of all purified TRIM28 PHD-bromodomain binders demonstrated that KI-T28-03 stabilizes TRIM28, whereas other compounds had weaker or destabilizing effects (**Supplementary Figure 17A-B**). Isothermal titration calorimetry (ITC) confirmed low micromolar binding of KI-T28-03 to full-length human TRIM28 protein (**Supplementary Figure 18A-B**). Computational modeling predicted high-confidence binding pockets for KI-T28-03 within the TRIM28 bromodomain **(Figure 5D)**, and structural modeling and sequence alignment showed that KI-T28-03 contacts poorly conserved residues within the bromodomain, consistent with selectivity **(Supplementary Figure 18C-D)**. To validate the predicted binding interface, we introduced a point mutation at F772 in the TRIM28 bromodomain (F772A), a residue predicted by structural modeling to make critical contact with KI-T28-03. Thermal-shift profiling showed that KI-T28-03 stabilizes wild-type TRIM28 but destabilizes the F772A mutant, confirming that F772 is required for productive compound engagement **(Supplementary Figure 18E)**. Dose-response assays revealed a half-maximal inhibitory concentration (IC_50_) of 4.85 µM in human MOLM13 leukemia cells **(Figure 5E)**, with similar low micromolar IC_50_ values in additional AML and B/Myelomonocytic leukemia cell lines, and substantially reduced potency in induced granulocyte– monocyte progenitors **(Supplementary Figure 19A-D)**. KI-T28-03 did not impair colony-forming capacity of human CD34^+^ progenitor cells **(Supplementary Figure 19E)**. Together, these data establish KI-T28-03 as a selective, low-micromolar, first-in-class TRIM28 inhibitor that both impairs leukemia cell proliferation and drives myeloid differentiation.

**Figure 5.**
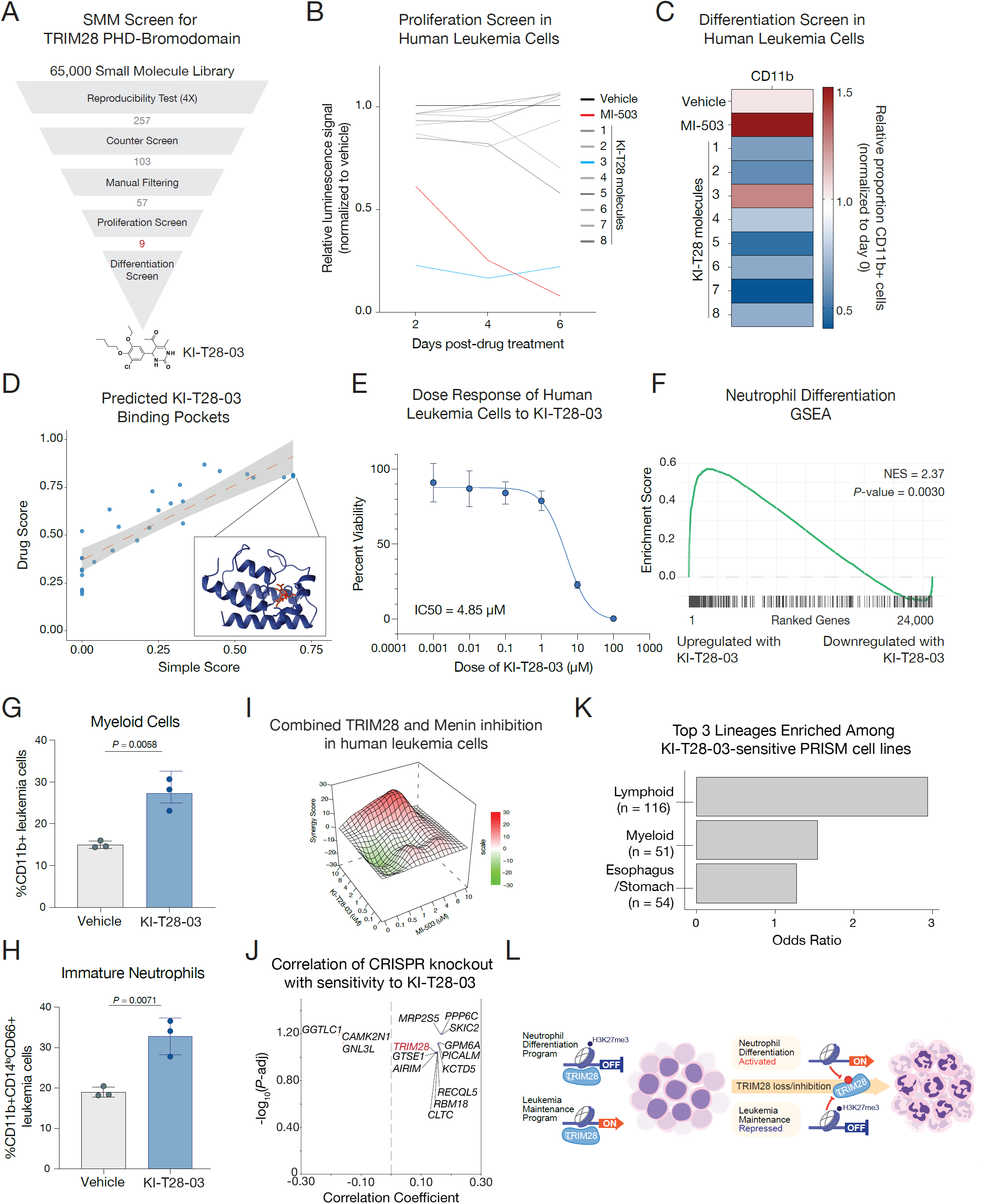
First-in-class TRIM28 inhibitor suppresses leukemia growth and drives neutrophil differentiation. **(A)** Schematic of the screening strategy to identify candidate small-molecule inhibitors of the TRIM28 PHD-bromodomain. **(B)** Proliferation curves for MOLM13 human leukemia cells treated with the indicated small-molecule inhibitors; cell viability was measured every 2 days for 6 days, normalized to DMSO, and mean viability for each compound was plotted (n = 3 replicates). Menin-MLL inhibitor (MI-503) was included as a positive control. **(C)** CD11b cell surface staining in MOLM13 cells treated with the indicated compounds for 6 days (n = 3 replicates). The average percentage of CD11b^+^ cells, normalized to DMSO, is shown as a heatmap. Menin-MLL inhibitor (MI-503) was included as a positive control. **(D)** Scatter plot of SimpleScore (*x*-axis) vs. DrugScore (*y*-axis) for predicted KI-T28-03 binding pockets on full-length TRIM28; the highest-scoring pockets are highlighted, with an inset showing the AlphaFold 3 model of the inhibitor bound to the top-ranking pocket (TRIM28 in purple, KI-T28-03 in red). **(E)** Dose-response curve for MOLM13 cells treated with six concentrations of KI-T28-03; viability was measured across replicates (n = 3), normalized to DMSO, and fitted with a nonlinear regression, with IC_50_ indicated in the bottom left. **(F)** GSEA plot showing enrichment of the bone marrow neutrophil gene signature (M39203) after 4 days of KI-T28-03 treatment in MOLM13 cells; NES and *P*-value shown. **(G)** Proportion of CD11b^+^ MOLM13 cells after KI-T28-03 treatment at its IC_50_ for 6 days (n= 3 replicates). **(H)** Proportion of immature neutrophils (CD11b^+^, CD14^lo^, CD66b^+^) in MOLM13 cells after KI-T28-03 treatment at its IC_50_ for 6 days (n = 3 replicates). **(I)** Topology map of ZIP synergy scores for combined KI-T28-03 and Menin-MLL inhibitor (MI-503) treatment in MOLM13 cells. **(J)** Scatter plot of the correlation between CRISPR knockout sensitivity (DepMap gene effect score; *x*-axis) and KI-T28-03 treatment sensitivity for each gene, with −log_10_(*P*-value) on the *y*-axis. Genes with significant positive correlation are highlighted in purple. **(K)** Enrichment scores (odds ratios) for each cancer lineage among the top 25% most KI-T28-03-sensitive cell lines. **(L)** Model summarizing the role of TRIM28 in leukemia maintenance and neutrophil differentiation, and the impact of TRIM28 inhibition.

To define the cellular programs engaged by KI-T28-03, we profiled drug-treated leukemia cells by RNA-Seq **(Supplementary Figure 20A)**. TRIM28 inhibition by KI-T28-03 significantly upregulated a neutrophil differentiation gene signature, as shown by gene set enrichment analysis **(Figure 5F, Supplementary Figure 20B)**, phenocopying our genetic *TRIM28* loss-of-function studies **(Figure 3B-C)**. We observed no significant changes in transposable element expression **(Supplementary Figure 20C)**. Consistent with these transcriptional changes, KI-T28-03 increased the proportion of CD11b^+^ myeloid leukemia cells and expanded an immature neutrophil population (CD11b^+^CD14^lo^CD66b^+^) **(Figure 5G-H)**. Subcellular fractionation and immunoblotting confirmed that TRIM28 remained predominantly nuclear and chromatin-associated following KI-T28-03 treatment **(Supplementary Figure 20D)**, indicating that KI-T28-03 disrupts TRIM28-dependent programs without globally displacing it from chromatin. Thus, pharmacologic TRIM28 inhibition phenocopies genetic *TRIM28* loss-of-function by releasing a differentiation block and driving leukemia cells toward a neutrophil-like fate.

We next evaluated the breadth of KI-T28-03 activity and its relationship to TRIM28 dependence. In human leukemia cells, combined treatment with KI-T28-03 and the Menin-MLL inhibitor MI-503^21^ produced strong synergistic growth inhibition across multiple dose combinations **(Figure 5I)**. Using PRISM profiling of 917 cancer cell lines^68^ **(Supplementary Figure 21A)**, we observed that sensitivity to KI-T28-03 strongly correlated with decreased viability upon CRISPR-mediated *TRIM28* knockout in DepMap^26^, but not with knockout of most other genes **(Figure 5J)**, supporting on-target activity. The same PRISM analysis identified 121 lines with KI-T28-03 IC_50_ values <10 µM, with the top 25% of sensitive lines most enriched for lymphoid, myeloid, and esophageal/stomach cancers **(Figure 5K, Supplementary Figure 21B)**. The preferential sensitivity of acute lymphoid and myeloid leukemias to KI-T28-03 is highly consistent with our initial functional genomics findings and Broad DepMap analyses of *TRIM28* dependence **(Figure 1B-C)**. These findings, together with our mechanistic data, support a model in which TRIM28 maintains a leukemia self-renewal program and suppresses neutrophil differentiation, and in which its loss or inhibition shifts leukemia cells toward a terminal neutrophil-like fate **(Figure 5L)**.

## Discussion

Functional genomic screening and mechanistic studies identify TRIM28 as a context-specific dependency in AML that sustains the leukemic state. TRIM28 acts as a scaffold for leukemia-specific complexes and enforces repression of myeloid differentiation programs, with a pronounced block in neutrophil lineage commitment. Genetic ablation of *TRIM28* drives leukemia cells into a neutrophil-like state with reduced proliferative potential and acquisition of terminal differentiation features. Guided by these insights, biochemical and cellular screens were used to develop a first-in-class small-molecule TRIM28 inhibitor that phenocopies key aspects of genetic *TRIM28* loss and shows selective activity in leukemia models. These findings define TRIM28 as a disease-relevant chromatin adaptor that governs leukemia cell fate and nominate both genetic *TRIM28* disruption and pharmacologic TRIM28 inhibition as therapeutic strategies in AML.

Mechanistically, TRIM28 sustains leukemic transcriptional programs through pathways that diverge from its canonical roles in heterochromatin assembly^32^ and retroelement silencing^47,69^. Instead, its PHD-bromodomain scaffolds context-dependent chromatin-modifying complexes that repress neutrophil-associated genes via H3K9me3-independent mechanisms, with associated dynamic remodeling of H3K27me3. Structural and biochemical characterization of the TRIM28 inhibitor KI-T28-03, which binds the PHD-bromodomain with low micromolar potency, demonstrates that this domain is a tractable pharmacologic entry point for modulating TRIM28 function. TRIM28 has also been implicated as a transcriptional regulator in breast cancer through interactions with EZH2 and SWI/SNF^70^, suggesting mechanistic parallels across tumor types despite distinct chromatin contexts. Together, these data support a model in which TRIM28 functions not as a static repressor but as a versatile chromatin adaptor that executes context-dependent transcriptional programs amenable to small-molecule intervention.

Although the mechanistic work centers on AML, functional genomics, DepMap analyses, and PRISM profiling indicate that acute lymphoid malignancies (e.g., B-cell acute lymphoblastic leukemia (B-ALL)) are also highly sensitive to *TRIM28* perturbation and TRIM28 inhibition. One possibility is that TRIM28 generally reinforces the malignant lineage of origin by suppressing alternative lineage programs, such that *TRIM28* loss in AML derepresses neutrophil-like transcriptional circuits, whereas in B-ALL, TRIM28 inhibition might unlock repressed T-cell-associated programs and promote lineage redirection. More broadly, the TRIM28 inhibitor described here provides a chemical probe to dissect how TRIM28 enforces lineage fidelity across lymphoid and myeloid cancers, and testing these lineage-switching hypotheses in engineered and patient-derived models will be an important direction for future work.

The neutrophil-like cells that emerge after *TRIM28* knockdown or TRIM28 inhibition arise from genetically altered leukemia cells and are unlikely to be identical to normal neutrophils, but their consistent induction defines a therapeutically exploitable differentiation trajectory. Given the rapid turnover of normal neutrophils, redirecting leukemia cells toward terminal myeloid differentiation may represent a viable treatment strategy, and restricting therapy-induced differentiation to this short-lived lineage has been proposed to reduce relapse and achieve cures in AML models^56^. The demonstration that a first-in-class TRIM28 inhibitor can drive neutrophil-like differentiation while impairing leukemia cell fitness provides a concrete pharmacologic implementation of this concept, and the delineation of TRIM28 functional domains, interactors, and druggable pockets offers a blueprint for further optimization of TRIM28-directed therapies.

TRIM28 inhibition emerges as a rational partner for Menin-directed therapy. Menin is a critical scaffold for KMT2A/MLL1 fusion-driven transcription^19,71^ and the target of clinically approved Menin-MLL inhibitors in high-risk leukemias^9^. However, *MEN1* mutations arising at relapse^5^ indicate that Menin inhibition alone may not provide durable control in all patients. Menin primarily sustains HOXA/MEIS1-like stemness programs^19,71^, whereas TRIM28 enforces repression of myeloid and neutrophil differentiation programs, indicating largely non-overlapping transcriptional modules. This division of labor, together with the strong synergy observed between TRIM28 inhibition and a Menin-MLL inhibitor (MI-503), suggests that TRIM28 targeting provides an orthogonal route to intercept leukemic self-renewal, even in settings of Menin-inhibitor resistance. Although formal *in vivo* and clinical studies are still required, the development of a first-in-class TRIM28 inhibitor and its cooperative activity with Menin inhibition raise the possibility that dual targeting of these chromatin adaptors could achieve more effective and durable responses than either agent alone.

## Limitations of study

This study defines TRIM28 as a leukemia-selective chromatin adaptor dependency and establishes a first-in-class small-molecule inhibitor that promotes neutrophil-like differentiation in acute leukemia, but several limitations should be considered.

First, although we employ multiple murine and human leukemia models, these systems cannot fully recapitulate the genetic and microenvironmental heterogeneity of patient AML, and the generalizability of TRIM28 dependence across additional subtypes remains to be established. Second, while our mechanistic data support a model in which the TRIM28 PHD-bromodomain scaffolds PRC2 and other chromatin regulators to repress neutrophil lineage genes, the full composition, dynamics, and locus specificity of these complexes in primary leukemias are not yet completely defined. Third, KI-T28-03 is a first-in-class small-molecule inhibitor with low micromolar potency, and we have not yet optimized its pharmacokinetic properties or evaluated its efficacy *in vivo*, so its direct translational potential remains to be determined. Finally, because we focus on leukemia, future work will be required to assess whether TRIM28-directed differentiation therapy is effective and tolerated in other TRIM28-dependent malignancies and in combination with additional standard-of-care agents.

## Acknowledgements

We would like to acknowledge all the members of the Soto-Feliciano, Lowe, and Koehler labs for their help and support in this project. We would like to thank Jacqueline Lees and Emma Montgomery for feedback on the manuscript. Colin Fowler, Tobias Coombs, David Chen (Chun-Wei Chen), and the Sànchez-Rivera Lab for experimental and reagent support. Alexey Soshnev for assistance with the model figure. This work was completed with assistance from the following core facilities at MIT, Koch Institute, and MSKCC: Swanson Biotechnology Center Flow Cytometry Core Facility (Glenn Paradis and Michele Griffin), Whitehead Institute Quantitative Proteomics Core (Fabian Schulte), Barbara K. Ostrom (1978) Bioinformatics & Computing Facility (Charlie Whittaker), BioMicro Center (Stuart Levine), and Preclinical Modeling (Aurora Burds-Connor).

- Blood Cancer United (formerly Leukemia and Lymphoma Society) SCOR (Y.M.S-F., S.M.A., I.C.)
- NIH/NIGMS R00 Award (R00-GM140265) (Y.M.S-F., S.M.A., A.A-O., O.S., I.C.)
- Ludwig Center at MIT (Y.M.S-F., S.M.A., Y.X., A.A-O., O.S., I.C.)
- NIH/NCI Core Grant (P30) (Y.M.S-F., SMA, A.A-O., O.S., I.C., G.J., F.J.S.R.)
- AACR Gertrude B. Elion Cancer Research Award (Y.M.S-F.)
- V Foundation V Scholar Award (Y.M.S-F.)
- MIT School of Science Dean’s Fellowship (2664134) (S.M.A.)
- HHMI Gilliam Fellowship (GT15758) (T.J.G.R.)
- NYU NIH Training Grant in Cell Biology (T32GM136542) (T.J.G.R.)
- Howard Hughes Medical Institute (S.W.L., M.P., F.J.S.R)
- Geoffrey Beene Chair for Cancer Biology (S.W.L., C-C.C., Y.J.H.)
- AML SPORE (S.W.L., V.N.)

## Author Contributions

S.M.A., C-C.C, S.W.L., and Y.M.S-F. conceived the study and wrote the manuscript with input from all the authors; S.M.A., C-C.C., Y.X., S.G., K.M., V.N., C.G., A.A-O, O.S., I.C., N.O., T.D., and Y.M.S-F. conducted experiments; S.M.A., C-C.C., T.J.G.R. and Y.J.H. analyzed RNA-Seq, ChIP-Seq, CUT&RUN, CRISPR screens data; S.M.A. and G.J. analyzed TRIM28 saturation mutagenesis screen data. S.M.A. and T.J.G.R. analyzed co-IP/MS data and DepMap co-dependency associations. Z.Y. performed AlphaFold 3 modeling. Y.M.S-F. and F.J.S.R. designed and cloned CRISPR libraries. Y.M.S-F, S.W.L., M.P., A.N.K., and K.V.R. supervised the study and secured funding.

## Materials and Methods

### Plasmids and Cloning

To generate stable Cas9-expressing cell lines, we used lentiCas9-Blast (Addgene, 52962). Mouse full-length or mutant Trim28 cDNA was cloned from a gene fragment (Twist Bioscience) into pCDH-EF1α-MCS-IRES-RFP (System Biosciences, CD531A-2). To express sgRNAs, we cloned these into the pUSEPR (hU6-sgRNA-EF1α-Puro-P2A-TurboRFP) lentiviral vectors^4^, sequences listed in **Supplementary Table 1A**. For sgRNA cloning, pUSEPR vectors were linearized with BsmBI (NEB) and ligated with BsmBI-compatible annealed and phosphorylated oligos encoding sgRNAs using T4 DNA ligase (NEB). All sgRNA sequences used in this study were generated using the Broad Institute CRISPick tool^72^ and are listed in **Supplementary Table 1B**.

To generate doxycycline-inducible shRNA cell lines, shRNAs were designed using the MSKCC Lowe Lab splashRNA tool^73^ and were cloned into miRE-LT3GEPIR plasmid (Addgene, 111177). All shRNA sequences are listed in **Supplementary Table 1C**. Briefly, 97mer oligos were PCR amplified and restriction sites were added to each oligo. Both the vector and shRNA oligo were digested with EcoRI and XhoI (NEB). Fragments were then purified using Gel Extraction and PCR Purification kits (Qiagen) and ligated using T4 DNA Ligase (NEB).

To recombinantly express and purify full-length human TRIM28 protein, the expression vector for FLAG-tagged TRIM28 (Clone ID: OHu28114, Accession Version: NM_005762.3) was purchased from GenScript.

### Lentiviral Transduction

Lentiviruses were produced by co-transfection of HEK293T cells (ATCC) with pUSEPR-EpiV2.0 sgRNA library **(Supplementary Table 1D)**, human TRIM28 tiling library **(Supplementary Table 1E)**, individual sgRNA plasmids, individual shRNA plasmids, mouse Trim28 cDNA or lentiCas9-Blast, and packaging vectors psPAX2 (Addgene, 12260), and pMD2.G (Addgene, 12259) using Lipofectamine2000 Reagent (Thermo Fisher). Viral supernatants were collected in cell culture media at 48- and 72-hours post-transfection, 0.45 µm-filtered and stored at -80 °C.

### Cell Culture

Mouse MLL-AF9 leukemia cells were kindly shared by David Chen (ChunWei Chen) (City of Hope) and were originally generated by transformation of female C57/BL6 mouse bone marrow Lin^-^Sca1^+^cKit^+^ (LSK) cells with an MSCV-IRES-GFP (pMIG) retrovirus expressing the human *MLL-AF9* fusion transgene, and transplanted into sub-lethally irradiated recipient mice as described previously^17,74^. Leukemic blasts were harvested from moribund mice and cultured *in vitro* in IMDM (Gibco) supplemented with 15% FBS (Gibco), mouse IL-6 (10 ng/µL, PeproTech), mouse IL-3 (10 ng/µL, PeproTech), mouse SCF (20 ng/µL, PeproTech), penicillin (100 U/mL, Gibco), streptomycin (100 µg/mL, Gibco), L-glutamine (2 mM, Gibco), and plasmocin (5 µg/mL, InvivoGen). Human leukemia cells MOLM13 (ATCC) were cultured in RPMI-1640 (Gibco) supplemented with 10% FBS (Gibco), penicillin (100 U/mL, Gibco), streptomycin (100 µg/mL, Gibco), L-glutamine (2 mM, Gibco), and plasmocin (5 µg/mL, InvivoGen). Human HEK293T (ATCC) cells were maintained in DMEM (Corning) supplemented with 10% FBS (Gibco), penicillin (100 U/mL, Gibco), streptomycin (100 µg/mL, Gibco) and plasmocin (5 µg/mL, InvivoGen). Cas9-expressing cells were generated by lentiviral transduction of lentiCas9-Blast followed by blasticidin (InvivoGen) selection and validation of Cas9 expression and activity. shRNA-expressing cells were selected with puromycin (Gibco) after recovery and before subsequent experiments. shRNA-expression was induced using 1-2 µg/mL of doxycycline (Millipore-Sigma) in the appropriate cell culture medium and maintained at the same concentration throughout the course of the experiment, unless otherwise noted. All cells were confirmed to be free of mycoplasma contamination and cultured at 37 °C and 5% CO_2_.

### Immunoblotting

Cell pellets were collected and washed with cold 1X PBS, spun down at 600xg for 5 minutes at 4 °C, and flash-frozen down at -80 °C. Pellets were lysed using 2X Laemmli buffer supplemented with 5% (w/v) β-mercaptoethanol (Thermo Fisher). Whole cell lysates were separated by SDS-PAGE (Thermo Fisher), transferred to a 0.45 µm PVDF membrane (Millipore), blocked in 5% non-fat milk in 1X TBS (Boston Bioproducts) plus 0.5% (w/v) Tween-20 (Sigma-Aldrich), probed with the primary antibodies (anti-TRIM28 (Bethyl, A300-275A), anti-Cas9 (CST, 14697S), anti-HA (CST, 3724S)), and detected with HRP-conjugated secondary antibodies (anti-rabbit (CST, 7074V) or anti-mouse (CST, 7076S)).

### Cell Surface Staining

Immunophenotyping of leukemia cells treated with Menin-MLL inhibitor (MI-503) (MedChem Express) or vehicle (DMSO) (Sigma-Aldrich), and/or after Trim28/TRIM28 knockdown, was done by collecting cells post treatment and staining using the indicated conjugated primary antibodies. Briefly, 1 million cells were collected per condition. Cell pellets were washed twice with Cell Staining Buffer (Biolegend, 420201), then samples were incubated with conjugated primary antibodies (1:100 dilution) for 30 minutes on ice and protected from light. Following primary antibody incubation, cells were washed twice with Cell Staining Buffer. Stained samples were analyzed on the LSRFortessa flow cytometer (BD Biosciences) and compensation was performed when appropriate. Data analysis was performed using FlowJo software (BD Biosciences). Fluorescently conjugated primary antibodies used were: PerCP anti-human CD14 (Biolegend, 325632), PE anti-human CD24 (Biolegend, 311106), Alexa Fluor 700 anti-human CD66b (Biolegend, 305114), PE/Cy7 anti-human CD68 (Biolegend, 333816), Alexa Fluor 647 anti-mouse/human CD11b (Biolegend, 101218), Brilliant Violet 421 anti-human CCR5 (Biolegend, 359118).

### Intracellular Staining

Detection of intracellular antigens was done using the Foxp3/Transcription Factor Staining Buffer Set (eBioscience) following the manufacturer’s guidelines. Briefly, 1 million cells were collected per reaction and fixed for 1 hour at room temperature and protected from light. Cell pellets were then washed twice before incubation with primary antibodies (1:100 dilution) for 1 hour at room temperature. Cell pellets were washed twice before incubation with fluorescently conjugated secondary antibodies (1:500 dilution) for 30 minutes at room temperature. Stained samples were analyzed on an LSR-Fortessa flow cytometer (BD Biosciences). Data analysis was performed using FlowJo software (BD Biosciences).

### Apoptosis Experiments

Cell death via apoptosis measurements were performed using Pacific Blue Annexin V Apoptosis Detection Kit with PI (Biolegend, 640928), according to the manufacturer’s instructions. Stained samples were analyzed on an LSRFortessa (BD Biosciences) flow cytometer. Data analysis was performed using FlowJo software (BD Biosciences).

### Reactive Oxygen Species Assay

Measurement of reactive oxygen species (ROS) in cells was adopted from Yokoyama et al.^58^. Cells were stimulated with 50 ng/mL of recombinant human G-CSF (Thermo Fisher) in reaction medium (HBSS + 0.5% BSA) for 15 minutes at 37 °C. Following stimulation, cells were incubated with 29 mM dihydrorhodamine 123 (DHR) (Sigma-Aldrich) in the reaction medium for 15 minutes at 37 °C. Cells were then washed twice and oxidized DHR was measured on an LSRFortessa (BD Biosciences) flow cytometer.

### *In vitro* Growth Competition Experiments

Cas9-expressing cells were virally transduced with the designated constructs (pUSEPR-sgRNA, pCDH-cDNA-RFP) in 12-well plates at ∼30% infection rate (3 infection replicates per experiment). Cells were monitored by flow cytometry over time using an LSRCelesta or LSRFortessa (BD Biosciences) flow cytometer and relative growth of sgRNA-containing cells was assessed. Flow cytometry data was analyzed with FlowJo software (BD Biosciences). The percentage of RFP^+^ cells was normalized to their respective T0 time-point values (assessed on day 2 post-transduction, as indicated in the figure legend). In shRNA-mediated knockdown experiments, cells were induced with 1 µg/mL of doxycycline (Millipore-Sigma). The proportion of GFP+ cells was monitored by flow cytometry over time using an LSRCelesta or LSRFortessa (BD Biosciences) flow cytometer.

### Chromatin-Focused and TRIM28 Saturation Mutagenesis CRISPR Screens

To ensure that most cells had a single sgRNA integration event, we determined the volume of viral supernatant that would achieve an MOI of ∼0.3 upon infection of a population of Cas9-expressing leukemia cells. Briefly, cells were plated at a concentration of 2.5 x 10^5^ per well in 12-well plates along with increasing volumes of library pool viral supernatant and polybrene (10 µg/mL, EMD Millipore). Cells were then centrifuged at 1500 rpm for 2 hours at 37 °C and incubated at 37 °C overnight. Viral infection efficiency was determined by the percentage of RFP+ cells assessed by flow cytometry on an LSRFortessa (BD Biosciences) instrument 72 hours post infection.

Each step of the screen, from infection to sequencing, was optimized to achieve a minimum representation of 1000X. 24 hours after infection, cells were pooled into T225 flasks (Corning) per infection replicate and selected with 2.5 µg/mL puromycin (Gibco) for 4 days. Subsequently, 6 million puromycin-selected cells were pelleted and stored at -20 °C (T0/Input population) while the rest were plated and cultured until the population reached ∼12 cumulative population doublings (Tf/Final population). Genomic DNA from screen cells was isolated using the DNeasy Blood & Tissue Kit (Qiagen) following the manufacturer’s guidelines.

As previously published^75^, we assumed that each cell contains approximately 6.6 pg of genomic DNA (gDNA). Therefore, deconvolution of the screen at 1000X required sampling ∼4 million (Chromatin-focused screen) or ∼ 1 million (TRIM28 tiling screen) x 6.6 pg of gDNA. We employed a modified 2-step PCR version of the protocol published by Doench et al.^76^. Briefly, we perform an initial ‘sgRNA-enrichment PCR’, whereby the integrated sgRNA cassettes are amplified from gDNA, followed by a second PCR to append Illumina sequencing adapters on the 5’- and 3’-ends of the amplicon, as well as a random nucleotide stagger and unique demultiplexing barcode on the 5’-end. Each ‘PCR1’ reaction contains 25 µL of Q5 High-Fidelity 2X Master Mix (NEB), 2.5 µL of PCR#1 Fwd Primer (10 µM), 2.5 µL of PCR#1 Rev Primer (10 µM), and 5 µg of gDNA in 20 µL of water (for a total volume of 50 µL per reaction). The number of ‘PCR1’ reactions was scaled accordingly. ‘PCR1’ amplicons were purified using the QIAquick PCR Purification Kit (Qiagen) and used as template for ‘PCR2’ reactions. Each ‘PCR2’ reaction contains 25 µL of Q5 High-Fidelity 2X Master Mix (NEB), 2.5 µL of a unique PCR#2 Fwd Primer (10 µM), 2.5 µL of PCR#2 Rev Primer (10 µM), and 300 ng of ‘PCR1’ product in 20 µL of water (for a total volume of 50 µL per reaction). We performed two ‘PCR2’ reactions per ‘PCR1’ product. Library amplicons were size selected on a 2.5% agarose gel, purified using the QIAquick Gel Extraction Kit (Qiagen), and sequenced on an NextSeq500 instrument (Illumina) (75nt single end reads). All primer sequences are available in Supplementary Table 1C. PCR Program for ‘PCR1’ and ‘PCR2’: 1) 98 °C x 30 s; 2) 98 °C x 10 s; 3) 65 °C x 30 s; 4) 72 °C x 30 s; 5) Go to step 2 x 24 cycles; 6) 72 °C x 2 min; 7) 4 °C forever.

### CRISPR Screens Analysis

For analysis of enrichment/depletion of sgRNAs, we used the MAGeCK algorithm, with the T0 replicates designated as the control timepoint. We then filtered to exclude sgRNAs with a control count mean < 10 reads to reduce spuriously enriching sgRNA hits. The rolling average of the guides and downstream figures were generated using python.

The human TRIM28 tiling screen sequencing results were demultiplexed into separate FASTQ files based on the sample barcode. Next, using a custom analysis script77, we filtered reads with an average Phred quality score below 30, and identified sgRNAs based on the protospacer sequence. Sequences with matching protospacers were used to generate sgRNA count tables that were subsequently used for MAGeCK analysis (v.0.5.9) of sgRNA enrichment/depletion.

### RNA Isolation and RNA-Sequencing (RNA-Seq)

Cell pellets were homogenized using the QIAshredder column (Qiagen), then total RNA was isolated from cells using the RNeasy kit (Qiagen) and quantified using NanoDrop (Thermo Fisher). For RNA-Seq experiments, quality and quantification of extracted RNA was assessed using NanoDrop and Bioanalyzer (Agilent). For the mouse RNA-Seq experiment, sequencing libraries were prepared using the NEBNext Ultra II RNA Library Prep Kit (NEB), following the manufacturer’s instructions. Sequencing was done using the NextSeq500 (Illumina) to obtain > 20 million 75bp single-end reads per sample. For the human RNA-Seq experiment, poly(A) mRNA enrichment and library preparation was performed by Novogene.

### RNA-Seq Analysis

RNA-Seq data were analyzed by removing adaptor sequences using Trimmomatic v0.3678. RNA-Seq reads were then aligned to GRCm38.91 (mm10) with STAR 2.6.1a^79^, and transcript counts were quantified using featureCounts v2.0.680 to generate a raw count matrix. Differential gene expression analysis was performed using the DESeq2 package^81^ implemented in *R* (http://cran.r-project.org/). Principal component analysis (PCA) was performed using the DESeq2 package in *R*. The likelihood ratio test (LRT) in DESeq2 was employed to identify genes with significant time-dependent expression changes across different treatment conditions. The full model formula was “∼ shTrim28 + Time + shTrim28:Time”, and the reduced model was “∼ shTrim28 + Time”. Differentially expressed genes (DEGs) were determined by Benjamini-Hochberg adjusted *P*-value <0.05. For heatmap visualization of DEGs, samples were *z*-score normalized, *k*-means clustered, and plotted to show distinct patterns across the sample groups using the ‘pheatmap’ package in *R*.

To determine the combined effects of multiple shRNAs targeting *TRIM28*, the RNA-Seq data were analyzed using the Seq-N-Slide pipeline (https://github.com/igordot/sns), through rna-star followed by rna-stargroups-dge routes, with quality control assessment (MultiQC, python/cpu/v2.7.15) and adaptor trimming (Trimmomatic, v0.36)^78^. Briefly, reads were aligned to the corresponding *mm10*/*GRCm38* mouse or *hg38*/*GRCh38* human reference genome with a splice-aware (STAR, v2.7.3a) alignment using the single-end protocol for mouse and paired-end for human samples, followed by featureCounts (subread/v1.6.3) to generate an RNA counts table^79,80^. Counts were normalized and tested for differential mRNA expression using negative binomial generalized linear models as implemented by DESeq2, v1.40.1 P^81^. A time-interaction term was calculated with the model design defined as “∼ group * time”, where the group corresponds to sh*Trim28* in mouse model or sh*TRIM28* in human cells, and time corresponds to each timepoint collected within each experiment. log_2_(foldchanges) of contrasts (c^t^β/√c^t^Σc) were shrunken following lfc-Shrink(type=“ashr”) to stabilize genes with low or variable counts. Differential expression was assessed by PCA or unsupervised hierarchical clustering and visualized by a volcano plot and TPM expression heatmap. Differentially expressed genes (DEGs) were analyzed for gene set enrichment analysis (GSEA) with *R* packages: fgsea v1.26.0 and msigdbr v7.5.1 via the pre-ranked method with 10,000 permutations, based on the adaptive multi-level splitting Monte Carlo approach. All *P*-values were adjusted with the Benjamini-Hochberg method.

### Chromatin Immunoprecipitation-Sequencing (ChIP-Seq)

Cross-linking ChIP in mouse leukemia was performed with 20 million cells per immunoprecipitation. Cells were collected, washed once with ice-cold PBS, and flash-frozen. Cells were resuspended in ice-cold PBS and cross-linked using 1% paraformaldehyde (PFA) (Electron Microscopy Sciences) for 5 minutes at room temperature with gentle rotation. Unreacted PFA was quenched with glycine (final concentration 125 mM) for 5 minutes at room temperature with gentle rotation. Cells were washed once with ice-cold PBS and pelleted by centrifugation (800*g* for 5 minutes). To obtain a soluble chromatin extract, cells were resuspended in 1 mL of LB1 (50 mM HEPES pH 7.5, 140 mM NaCl, 1 mM EDTA, 10% glycerol, 0.5% NP-40, 0.25% Triton X-100, 1X complete protease inhibitor cocktail (Roche)) and incubated at 4 °C for 10 minutes while rotating. Samples were centrifuged (1400*g* for 5 minutes), resuspended in 1 mL of LB2 (10 mM Tris-HCl pH 8.0, 200 mM NaCl, 1 mM EDTA, 0.5 mM EGTA, 1X complete protease inhibitor cocktail (Roche)), and incubated at 4 °C for 10 minutes while rotating. Finally, samples were centrifuged (1400*g* for 5 minutes) and resuspended in 1 mL of LB3 (10 mM Tris-HCl pH 8.0, 100 mM NaCl, 1 mM EDTA, 0.5 mM EGTA, 0.1% sodium deoxycholate, 0.5% N-Lauroylsarcosine, 1X complete protease inhibitor cocktail (Roche)). Samples were homogenized by passing 7-8 times through a 28-gauge needle and Triton X-100 was added to a final concentration of 1%. Chromatin extracts were sonicated for 14 minutes using a E220 focused ultrasonicator (Covaris). Lysates were centrifuged at maximum speed for 10 minutes at 4 °C and 5% of supernatant was saved as input DNA. Beads were prepared by incubating them in 0.5% BSA in PBS and antibodies overnight (100 µL of Dynabeads Protein A or Protein G (Invitrogen) plus 20 µL of antibody). Antibodies used were: anti-TRIM28 (Bethyl, A300-275A) and anti-H3K9me3 (abcam, 8898). Antibody-Beads mixes were washed with 0.5% BSA in PBS and then added to the lysates overnight while rotating at 4 °C. Beads were then washed six times with RIPA buffer (50 mM HEPES pH 7.5, 500 mM LiCl, 1 mM EDTA, 0.7% sodium-deoxycholate, 1% NP-40) and once with TE-NaCl Buffer (10 mM Tris-HCl pH 8.0, 50 mM NaCl, 1 mM EDTA). Chromatin was eluted from beads in Elution Buffer (50 mM Tris-HCl pH 8.0, 10 mM EDTA, 1% SDS) by incubating at 65 °C for 30 minutes while shaking, supernatant was removed by centrifugation, and crosslinking was reversed by further incubating chromatin overnight at 65 °C. The eluted chromatin was then treated with RNaseA (10 mg/mL, Milli-pore-Sigma) for 1 hour at 37°C and with Proteinase K (Roche) for 2 hours at 55 °C. DNA was purified by using phenol-chloroform extraction followed with ethanol precipitation. The NEBNext Ultra II DNA Library Prep Kit (NEB) was used to prepare samples for sequencing on a NextSeq500 (Illumina) (75 bp read length, single-end).

### ChIP-Sequencing Data Analysis

ChIP-Seq reads were aligned using *R*subread’s align method and predicted fragment lengths calculated by the ChIPQC *R* Bioconductor package^82,83^. Normalized, fragment extended signal bigWigs were created using the rtracklayer *R* Bioconductor package^84^. Peak calls were made in MACS2 software^85^. Read counts in peaks were calculated using the featureCounts method in the Rsubread library^83^. Differential ChIP-Seq signals were identified using the binomTest from the edgeR *R* Bioconductor package^86^. Annotation of genomic regions to genes, biological functions, and pathways were performed using the ChIPseeker *R* Bioconductor package^87^. Metapeak plots were produced using the soGGi package and ChIP-Seq signal heatmaps generated using the Deeptools and profileplyr software^88^. Plots showing ChIP-Seq read signal over transcription start sites (TSSs) were made with the ngs.plot software package (v2.61)^89^. Overlaps between peak sets were determined using the ChIPpeakAnno *R* Bioconductor package with a maximum gap between peaks set to 1kb^90^. Peaks were annotated with both genes and the various types of genomic regions using the ChIPseeker R Bioconductor package^87^. Range-based heatmaps showing signal over genomic regions were generated using the soGGi and profileplyr R Bioconductor package to quantify read signal and group the peak ranges and the deepTools software package (v3.3.1) to generate the heatmaps^88^. Any regions included in the ENCODE blacklisted regions of the genome were excluded from all region-specific analyses^91^.

### CUT&RUN

Human AML cells (MOLM13) cells were initially plated at 300,000 cells/mL with 1 µg/mL doxycycline (Millipore-Sigma), cells were re-plated and doxycycline was re-applied after 48 hours. After 4 days, 500,000 leukemia cells were collected per CUT&RUN reaction. Nuclei were extracted using CUTANA Nuclei Extraction Buffer (Epicypher, 21-1026) according to the manufacturer’s instructions. CUT&RUN was subsequently performed using the CUTANA ChIC/CUT&RUN Kit (Epicypher, 14-1048). Extracted nuclei were conjugated to Concanavalin A beads and incubated with 2 µg of primary antibody overnight at 4 °C. Primary antibodies used were: anti-KAP1 (TRIM28) (abcam, 10484), anti-H3K9me3 (abcam, 8898), and anti-H3K27me3 (CST, 9733S). MNase conjugation and digestion were done the following day. CUT&RUN fragments were purified using SPRI beads and quantified using 1X DS DNA High Sensitivity Reagents using Qubit (Thermo Fisher). Library preparation was performed using CUTANA CUT&RUN Library Prep Kit (Epicypher, 14-1001) according to manufacturer’s instructions. Libraries were quantified using Tapestation (Agilent) and inspected for expected fragment size distribution.

### CUT&RUN Data Analysis

Sequencing was done using the NovaSeq X Plus (Illumina) to obtain >10 million 150bp paired-end reads at Novogene. FASTQ files were trimmed using TrimGalore^92^ and aligned using Bowtie2^93^. Peak calls were made in MACS2 software^85^. Annotation of genomic regions to genes, biological functions, and pathways were performed using the ChIPseeker *R* Bioconductor package^87^. CUT&RUN signal heatmaps generated using the Deeptools and profileplyr software^88^. Overlaps between peak sets were determined using the ChIPpeakAnno *R* Bioconductor package with a maximum gap between peaks set to 1kb^90^. Peaks were annotated with both genes and the various types of genomic regions using the ChIPseeker *R* Bioconductor package^87^. Range-based heatmaps showing signals over genomic regions were generated using the deepTools software package (v3.3.1)^88^. Any regions included in the ENCODE blacklisted regions of the genome^94^ were excluded from all region-specific analyses.

### Dependency Analysis using Broad Institute DepMap Database

The 808 cancer cell lines in the Broad Institute DepMap were segregated into leukemia models and non-leukemic models based on the description of the primary disease. The average dependency score for each gene was calculated among leukemic and non-leukemic cell lines. A leukemic-specific dependency was defined as a dependency score 150% (1.5 times) higher in leukemic models compared to non-leukemic. The results were visualized using Python.

### Functional Annotations of Gene Sets

Pathway enrichment analysis was performed using the indicated gene sets in enrichR^95^. Significance of the tests was assessed using combined score, described as c = log(p) * z, where c is the combined score, *P* is Fisher’s exact test *P*-value, and *z* is *z*-score for deviation from expected rank, as well as by adjusted *P*-values defined by enrichR.

### TRIM28 Co-Immunoprecipitation Followed by Quantitative Mass Spectrometry (Co-IP/MS)

20-million of human embryonic kidney cells (HEK293T) (ATCC) or 60-million of human AML cells (MOLM13) (ATCC) were collected as input for each biochemical pulldown. Cell pellets were then lysed using NETN Lysis Buffer (20 mM Tris-HCl pH8.0, 1 mM EDTA, 150 mM NaCl, 0.5% NP-40, 10% glycerol) completed with protease inhibitors (Roche), 1 mM DTT, and 1 mM PMSF. Sonication [5sec x 3 cycles] was then done using a Diagenode Pico sonication device to ensure lysate homogenization. Cell lysates were pre-cleared by incubating with a mixture of Protein A and Protein G beads for 30 minutes at 4 °C. Following pre-clearing, the lysates were incubated with Anti-HA magnetic beads (Thermo Scientific) for 30 minutes at room temperature. The beads were then washed 5 times with ice-cold NETN lysis buffer before being resuspended in water and sent for mass spectrometry. Samples were subjected to denaturation and in-solution digestion before Orbitrap liquid chromatography coupled to tandem mass spectrometry (Thermo Scientific) with 60-minute Data-Independent Acquisition (DIA) mode was performed at the Whitehead Institute Proteomics Facility. Protein abundance counts were normalized to respective whole cell lysates. Differential protein quantification was calculated using the eBayes (fit, trend = TRUE) as recommended by the DEqMS pipeline^96^. Results were visualized in *R* (v4.5.1). We performed gene set enrichment analysis (GSEA, fgsea v.1.35.697) ranking interactors by the signed enrichment score and used as input for pre-ranked enrichment testing against MSigDB gene sets (*R* package: msigdbr v.25.1.198).

### AlphaFold 3 Structural Modeling

The three-dimensional structure of human TRIM28 was predicted using AlphaFold 3 web server (https://alphafoldserver.com/), where default settings were used. Protein sequences were provided in FASTA format using UniProt canonical sequences. Truncated isoform sequences were adapted from human TRIM28 protein sequence (Q13263) according to the UniProt’s domain annotations. The predicted TM-score (pTM) is a global confidence metric produced by AlphaFold models to estimate the accuracy of the overall fold topology of a predicted structure. Values range from 0 to 1, with higher values indicating greater confidence in the predicted global fold. The figure displaying structural information was prepared and visualized using UCSF ChimeraX.

### DepMap TRIM28 Co-Dependency Analysis

To identify lineage-specific gene co-dependency networks, we analyzed CRISPR-Cas9 gene effect scores from the DepMap Public 25Q2 release (www.depmap.org), focusing on myeloid hematologic cancer models (n = 87). Dependency scores were retrieved from the CRISPRGeneDependency dataset and filtered to retain only cell lines annotated as myeloid based on Oncotree lineage classifications. Pairwise Pearson correlation coefficients were computed across all genes using the Hmisc::rcorr() function (v.5.2.399). The resulting gene-gene correlation matrix was converted into a long-format table containing the correlation coefficients, *P*-values, and sample sizes between each gene pair. To adjust for multiple hypothesis testing, *P*-values were corrected using the Benjamini-Hochberg (BH) method and resulting FDR values were used to compute signed enrichment scores (i.e., signed -log_10_(FDR)). We then extracted all gene pairs where TRIM28 was either the row or column gene, yielding a TRIM28-centric co-dependency profile. TRIM28 co-dependencies were ranked by signed enrichment scores to prioritize biologically meaningful associations. To gain functional insight into these dependencies, we performed gene set enrichment analysis (GSEA, fgsea v.1.35.697) using the ranked TRIM28 co-dependencies as input for pre-ranked enrichment testing against MSigDB gene sets (*R* package: msigdbr v.25.1.198). All data wrangling and visualization were performed in R (v.4.5.1).

### Small-Molecule Microarray Screening

Small-molecule microarrays (SMM) were prepared and analyzed against a recombinant human TRIM28 PHD-bromodomain protein (624-811aa), as previously described^67^. Approximately 10,000 printed features were present on each SMM slide, consisting of 5,000 unique compounds printed in duplicate. In total, 65,000 compounds were screened. Each slide was screened in duplicate, resulting in four replicate measurements per compound. Briefly, purified His6-tagged TRIM28 protein (Active Motif, 31441) was incubated on glass slides bearing printed compounds at a concentration of 1 µg/mL in TBS buffer supplemented with 0.5X BlockerTM FL Fluorescent Blocking Buffer (Thermo Fisher Scientific) for 1 hour at room temperature. Slides were washed twice with TBST buffer for 2 min each, and binding was detected using a DyLightTM 650-conjugated monoclonal anti-His6 antibody (Invitrogen, MA1-21315-D650) at a 1:1000 dilution in 0.5X Blocker FL Fluorescent Blocking Buffer. Slides were then subjected to three additional washes with TBST, followed by a brief rinse with TBS and ultrapure water. After drying by centrifugation, slides were immediately scanned using a SureScan microarray scanner (Agilent).

### Small-Molecule Microarray Screening Analysis

Scanned slide images were analyzed using GenePix Pro software (Molecular Devices) for spot alignment with the corresponding GenePix Array List (GAL) file. The resulting GPR files were processed using a data analysis workflow implemented in Pipeline Pilot (BIOVIA) and the KNIME Analytics Platform. Hit calls were determined as previously described^100,101^. The *Z*- score for each feature was calculated as below from signal-to-noise ratio (SNR) values using the mean and standard deviation across each subarray, as robust estimators resulted in unstable *Z*-values likely due to an extremely skewed SNR distribution with a very small median absolute deviation.

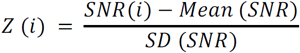

The SNR for an individual feature is denoted by SNR (i). The mean SNR value across all features in the subarray is represented by Mean (SNR). SD (SNR) indicates the standard deviation of SNR values for the entire set of features within the subarray. Assay positives were identified based on the following criteria: a Z-score > 3 and a SNR > 1.3 in all four replicates. Putative hits were further verified by manual inspection of slide images, and false positives were excluded through counter-screening assays.

### Small-Molecule Proliferation Experiments

MOLM13 cells were seeded at a density of 30,000 cells in 200 µL of appropriate media in a non-TC-treated 96 well plate. Cells were treated with 2 µL of small-molecule inhibitor resuspended in 10% DMSO or equal volume of vehicle. Cells were plated in triplicate for technical replicates. Cells were split and a small-molecule inhibitor was added every 2 days, and viability was measured using CellTiter-Glo (Promega) 4 or 6 days after initial plating. Viability was measured as luminescence and values were averaged across technical replicates and normalized to the vehicle at that day. Results were visualized using GraphPad Prism 10.

### Isothermal Titration Calorimetry (ITC)

ITC experiments were conducted at 25 °C using a MicroCal PEAQ-ITC system (Malvern Panalytical Ltd., Worcestershire, UK). The concentration of recombinantly purified human TRIM28 protein was measured by BCA. The KI-T28-03 was dissolved in the ITC buffer (25 mM HEPES buffer, 200 mM NaCl, and 5% DMSO) at 500 µM. Before the formal run, a deep soak cleaning procedure was performed according to the manufacturer’s instructions for optimal conditioning. The protein, after adding 5% DMSO for buffer matching, was loaded into the sample cell and then titrated with the small molecule. The volume loaded for the protein and small molecule are 300 µL and 70 µL, respectively, as recommended by the manufacturer’s instructions. For each titration experiment, the first injection was performed using µL followed by 18 additional injections at 2 µL per injection. The initial delay for titration is 60 seconds, while for later titrations, the spacing is 150 seconds, and each titration takes 4 seconds. The first injection was considered void and was automatically removed from the data analysis. The resulting heat plots were analyzed by using the MicroCal PEAQ-ITC Analysis software (Malvern Panalytical Ltd.).

### nano Differential Scanning Fluorimetry (nanoDSF)

nanoDSF to demonstrate thermal shift upon small molecules and protein binding was done with Prometheus Panta nanoDSF (Nanotemper Technologies). Small molecules including KI-T28-03 were dissolved in 100% DMSO to make 10 mM stock solutions which were diluted 1:10 in 100% DMSO to make 1 mM drug solutions. 50 µL of recombinant TRIM28-FLAG in nanoDSF assay buffers (25 mM HEPES, 250 mM NaCl, pH=7.5) at 0.4 mg/mL was mixed with 0.5 µL of 1 mM drug in DMSO to achieve a final 10 µM drug concentration in nanoDSF assay buffers with 1% DMSO, with nodrug DMSO control for each assay. This protein and drug mixture is incubated at room temperature for 20 minutes before transfer to nanoDSF Grade Standard Capillaries. Capillaries containing 10 µL protein drug mixture in duplicates were inserted into the machine, the temperature was increased at a rate of 1 °C/min from 25 °C to 95 °C, and the melting curve of each capillary was fitted by Prometheus Panta Analysis software to determine the T_m_. The protein’s thermal shift ΔT_m_ between DMSO control and drug duplicates were analyzed and reported.

### Recombinant Human TRIM28-FLAG Expression and Purification

HEK293T (ATCC) cells were cultured in Dulbecco’s Modified Eagle’s medium (DMEM) supplemented with 10% fetal bovine serum (FBS) with 1% penicillin/streptomycin and maintained in a tissue culture incubator at 37 °C with 5% CO_2_. 8 million cells were seeded per 15-cm dishes 16 hours before transfection in 20 mL DMEM with 10% FBS. Cells were transfected with PEI MAX® (Polyscience, 49553-93-7) according to the manufacturer’s instructions. 30 µg of TRIM28-FLAG plasmid in 0.5 mL Opti-MEM reduced serum medium (Gibco) and 60 µL of 1 mg/mL PEI Max in 0.5 mL of OptiMEM reduced serum medium were mixed rigorously to form the transfection mixture per plate. After 20 minutes of RT incubation, 1 mL of the mixture was added to HEK293T cells in each plate. After 48 hours of transfection, cells were harvested and centrifuged at 300g centrifugal force for 5 minutes. The pellet was lysed with 0.5 mL/plate of Mammalian Lysis Buffer (Promega, G6500) supplemented with PhosSTOP (Roche, 4906845001) and protease inhibitor cocktail (Roche, 11836170001) with gentle 30s vortexing. After incubating on ice for 40 minutes, the mixture was centrifuged at maximum speed (>18,000g) on tabletop centrifuges for 10 minutes at 4 °C. Anti-FLAG magnetic agarose (Thermo Fisher Scientific, A36797) was washed 3 times with Mammalian Lysis Buffer before adding the supernatant. The mixture of supernatant and pre-washed magnetic agarose was incubated on rotation at 250 rpm overnight at 4 °C. Beads were collected with a magnetic stand and washed twice with PBS and once with purified water. 100 µL of a PBS solution of 1.125 mg/mL 3X FLAG peptide (APExBIO, A6001) was added to the beads and incubated at 1,400 rpm for 10 minutes at room temperature. The supernatant was collected and the elution process was repeated one more time. The combined solution was added to Amicon Ultra0.5, 30 kDa (Merck-Millipore), and the buffer was adjusted to the following system (25 mM HEPES, 300 mM NaCl, 10% glycerol, 0.04% Triton X-100, mM DTT). Protein purity was determined by SDS-PAGE and can be used fresh for immediate assays or stored in -80 °C.

### Growth Inhibition Assay

MOLM13 cells were seeded at a density of 30,000 cells in 200 µL of media in a non-TC treated 96-well plate with 2 µL of increasing dilutions of small-molecule inhibitor or equal volume of vehicle. Cells were plated in triplicate for technical replicates. Cells were split and fresh inhibitor was reapplied after 48 hours. Viability was measured using CellTiter-Glo (Promega) after 4 days. Percent viability at each concentration was calculated by averaging across replicates and normalizing to the vehicle control. Results were visualized and IC_50_ values were calculated using GraphPad Prism 10.

### PRISM Screen

KI-T28-03 solution (10 mM in DMSO) was submitted to the Broad Institute PRISM project^68^. Briefly, 917 cancer cell lines were barcoded individually and pooled together for growth inhibition experiments. Cells were pooled together and incubated with increasing dilutions of KI-T28-03 for 5 days. Pools were then sequenced and deconvoluted to determine the viability of each cell line at each dose of KI-T28-03 inhibitor. Results were visualized and analyzed using *R* and Python.

### Statistical and Correlation Analyses

Statistical tests were used as indicated in figure legends. Generation of plots and statistical analyses were performed using Prism 10 (GraphPad). Error bars represent standard deviation, unless otherwise noted. We used Student’s *t*-test (unpaired, two-tailed) to assess significance between experimental and control groups and to calculate *P*-values. *P* < 0.05 was the threshold for statistical significance.

Gene expression measured by transcript counts was extracted from RNA-Seq experiments in human AML cells (MOLM13) and from human embryonic kidney cells (HEK293Ts) from Tchurikov *et al*.^102^. The log_2_(fold-change) expression in MOLM13 cells vs. HEK293T cells for each gene was calculated. This value was plotted against log_2_(fold-change) for TRIM28 interactome determined by co-IP/MS. The Pearson’s correlation coefficient was calculated. Results were visualized using *R* v4.3.2.

## Supplementary Tables

- Supplementary Table 1: Sequences of sgRNA (library and individual) and primers
- Supplementary Table 2: CRISPR screenings results
- Supplementary Table 3: ChIP-Seq/CUT&RUN results
- Supplementary Table 4: RNA-Seq results
- Supplementary Table 5: Quantitative proteomics results
- Supplementary Table 6: KI-T28-03 results

