## Supplementary Figures for "Epigenetic Reactivation of Lineage Differentiation to Target Leukemia"

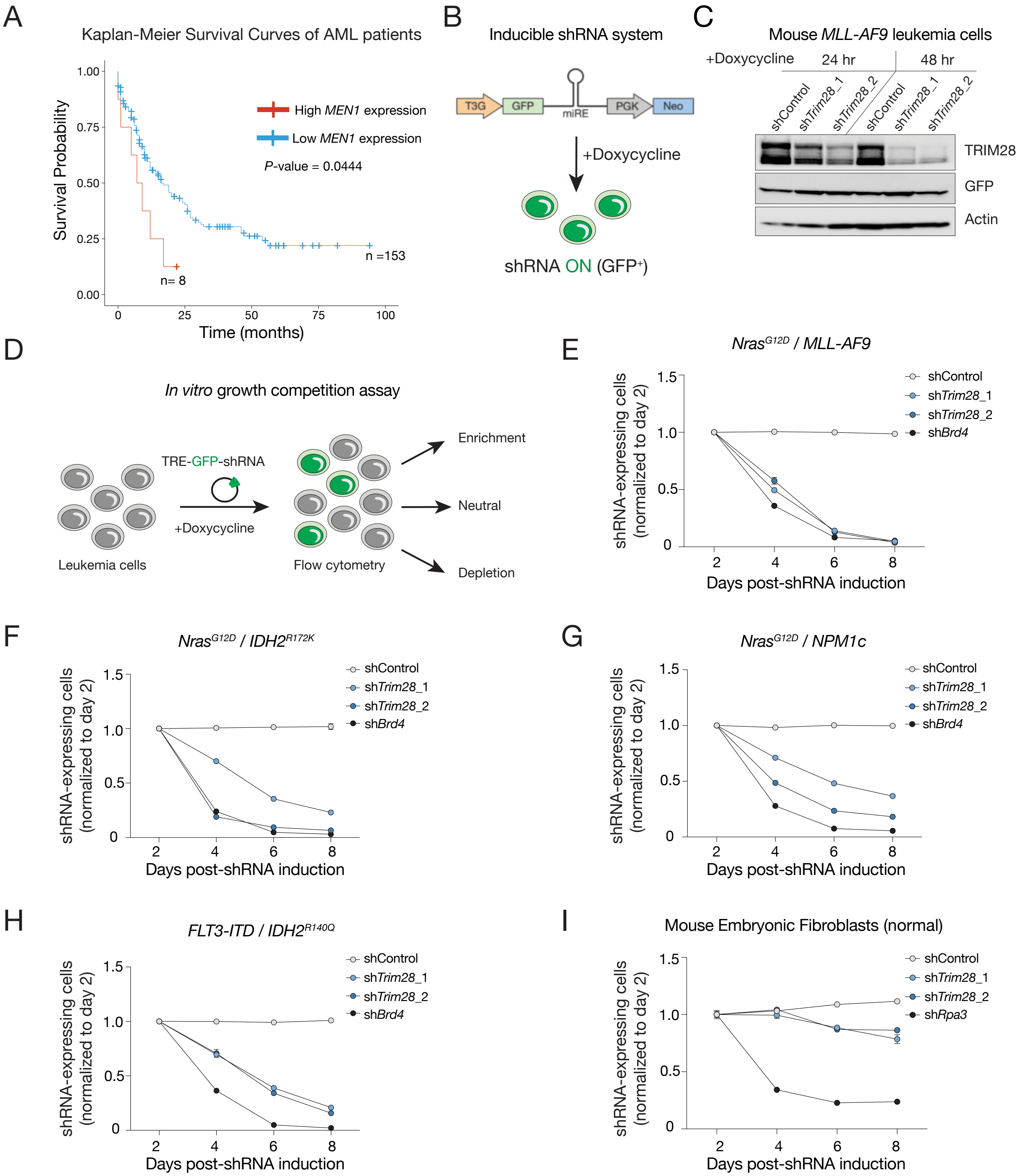


**Supplementary Figure 1 – *Trim28* is a leukemia-specific dependency in mouse models of acute myeloid leukemia. (A)** Survival data from TCGA AML patients stratified by high (n = 8) versus low (n = 153) *MEN1* expression; *P*-value calculated by log-rank test. **(B)** Schematic of the doxycycline-inducible shRNA construct. **(C)** Immunoblot of mouse MLL-AF9 cells engineered with doxycycline-inducible shRNAs (non-targeting shControl; *Trim28*-targeting sh*Trim28*_1 and sh*Trim28*_2) co-expressing *GFP*, before and 1 and 2 days after shRNA induction. **(D)** Schematic of doxycycline-inducible shRNA-mediated competition experiments. **(E-H)** Competition experiments in mouse leukemia cells with distinct driver mutations, indicated at the top of each panel, engineered with doxycycline-inducible shRNAs (non-targeting shControl; pan-essential control sh*Brd4*; *Trim28*-targeting sh*Trim28*_1 and sh*Trim28*_2) co-expressing *GFP*. Cells were treated with 1 µg/mL doxycycline (n = 2 replicates), and the proportion of GFP+ cells was quantified every 2 days over an 8-day period by flow cytometry. **(I)** Competition experiment in mouse embryonic fibroblasts (MEFs) engineered with doxycycline-inducible shRNAs, co-expressing *GFP*. Cells were treated with 1 µg/mL doxycycline (n = 3 replicates), and the proportion of GFP+ cells was quantified every 2 days over an 8-day period by flow cytometry.

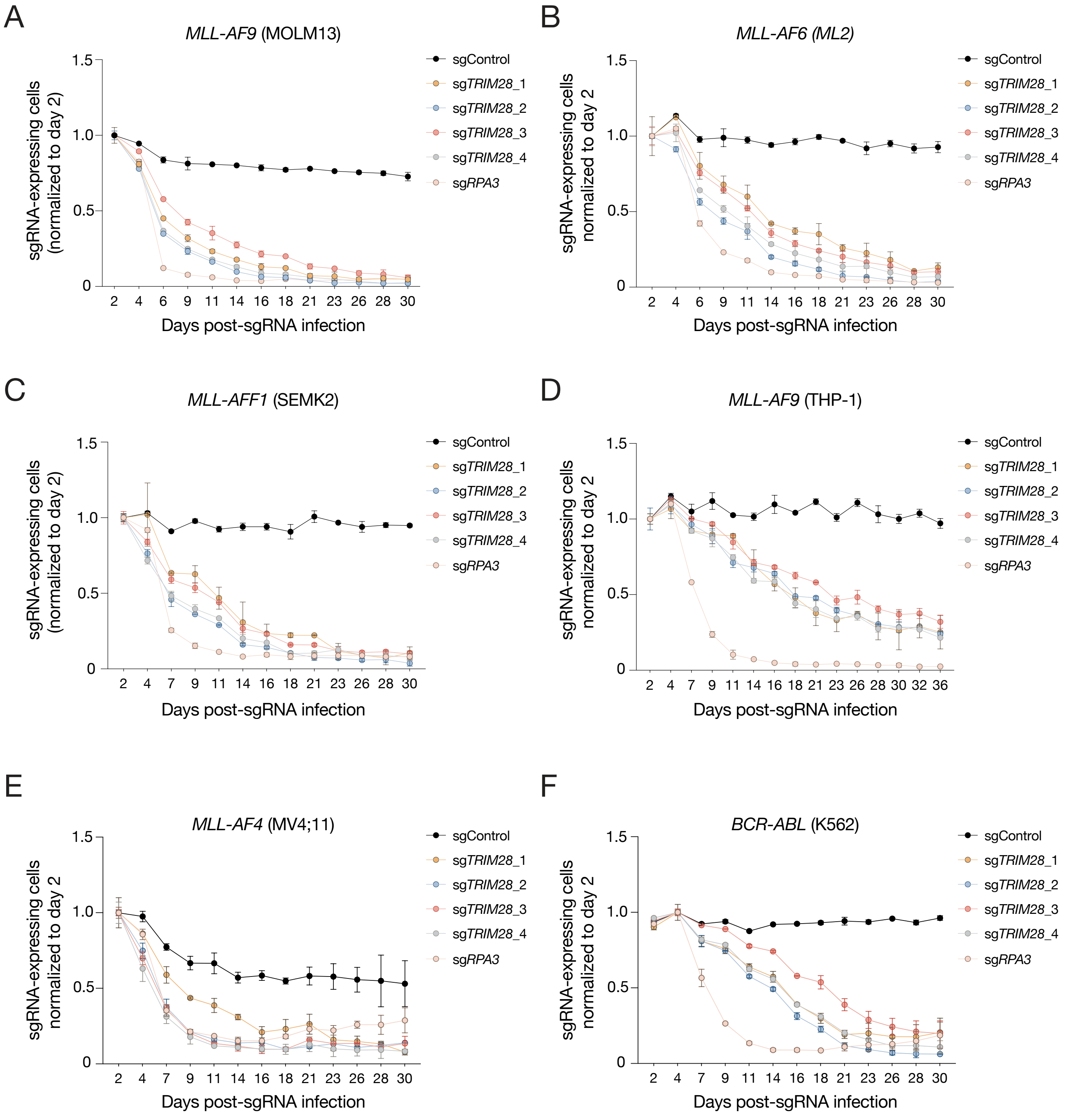


**Supplementary Figure 2 – Human leukemia cell lines are sensitive to *TRIM28* loss. (A-F)** Competition experiment in human leukemia cells with distinct driver mutations (indicated at the top of each panel; cell line name listed below), infected with sgRNAs (non-targeting sgControl; *TRIM28*-targeting sg*TRIM28*_1, sg*TRIM28*_2, sg*TRIM28*_3, sg*TRIM28*_4; and sg*RPA3*) co-expressing *GFP*, and the proportion of GFP⁺ cells was quantified every 2 days over a 30-day period using flow cytometry (n = 3 replicates).

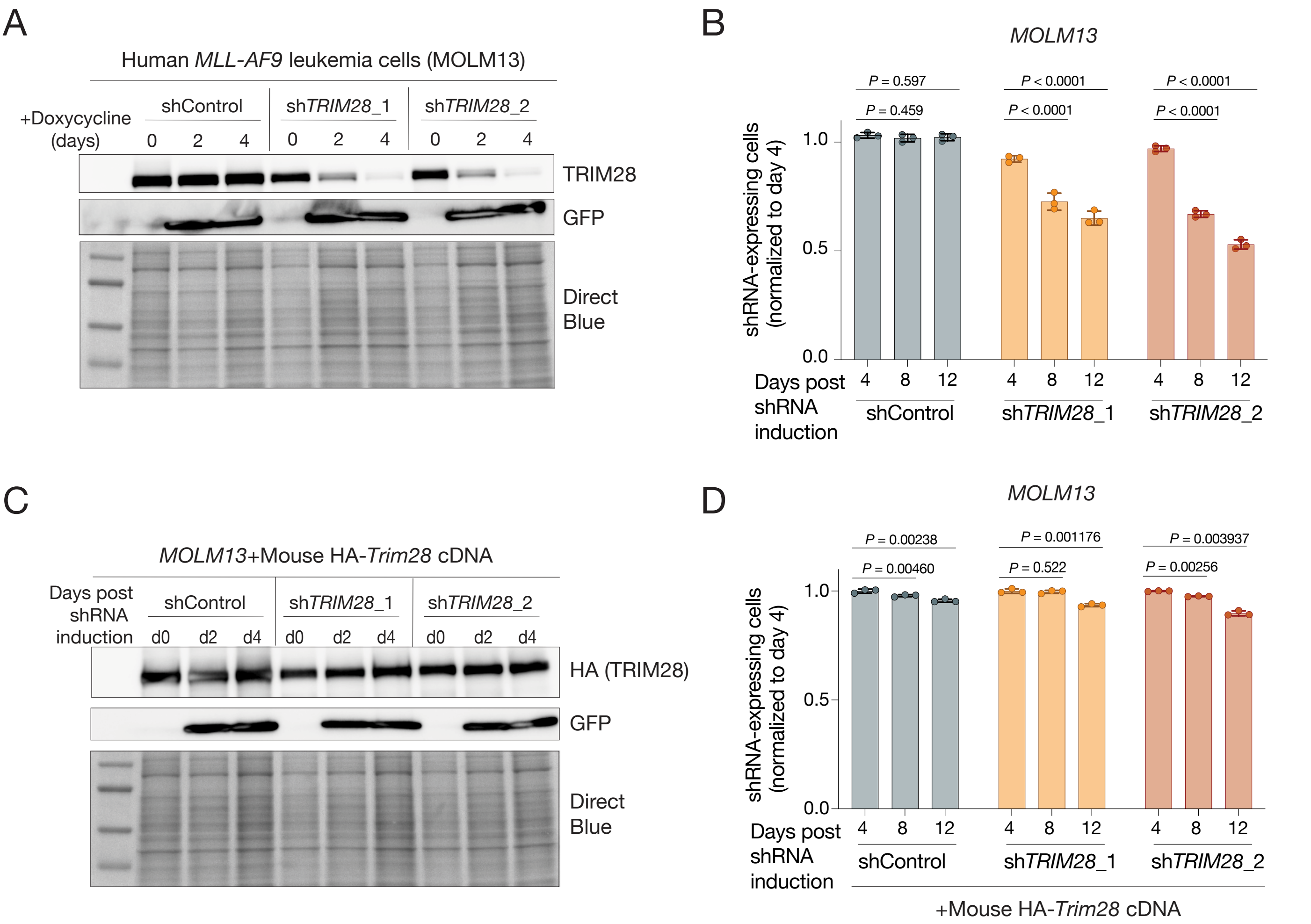


**Supplementary Figure 3 – *TRIM28* knockdown impairs proliferation of human leukemia cells and is rescued by mouse *Trim28* cDNA expression. (A)** Immunoblot of human leukemia cells (MOLM13) engineered with doxycycline-inducible shRNAs, co-expressing *GFP*. Cell pellets were collected before and 2 and 4 days after shRNA induction. The immunoblot shows TRIM28 (top), GFP (middle), and Direct Blue staining below as a loading control. **(B)** Competition experiment in MOLM13 cells engineered with doxycycline-inducible shRNAs, co-expressing *GFP*. Cells were treated with 1 µg/mL doxycycline and the proportion of $GFP^+$ cells was quantified every 4 days over a 12-day period by flow cytometry (n = 3 replicates). **(C)** Immunoblot of MOLM13 cells constitutively expressing HA-tagged full-length (FL) mouse *Trim28* cDNA, which is resistant to the human-targeting shRNAs, and additionally engineered with doxycycline-inducible shRNAs, co-expressing *GFP*. Cell pellets were collected before and 2 and 4 days after shRNA induction. The immunoblot shows HA as a marker for TRIM28 (top), GFP (middle), and Direct Blue staining below as a loading control. **(D)** Competition experiment in MOLM13 cells constitutively expressing HA-tagged FL-*Trim28* mouse cDNA and engineered with doxycycline-inducible shRNAs. Cells were treated with 1 µg/mL doxycycline and the proportion of $GFP^+$ cells was quantified every 4 days over a 12-day period by flow cytometry (n = 3 replicates).

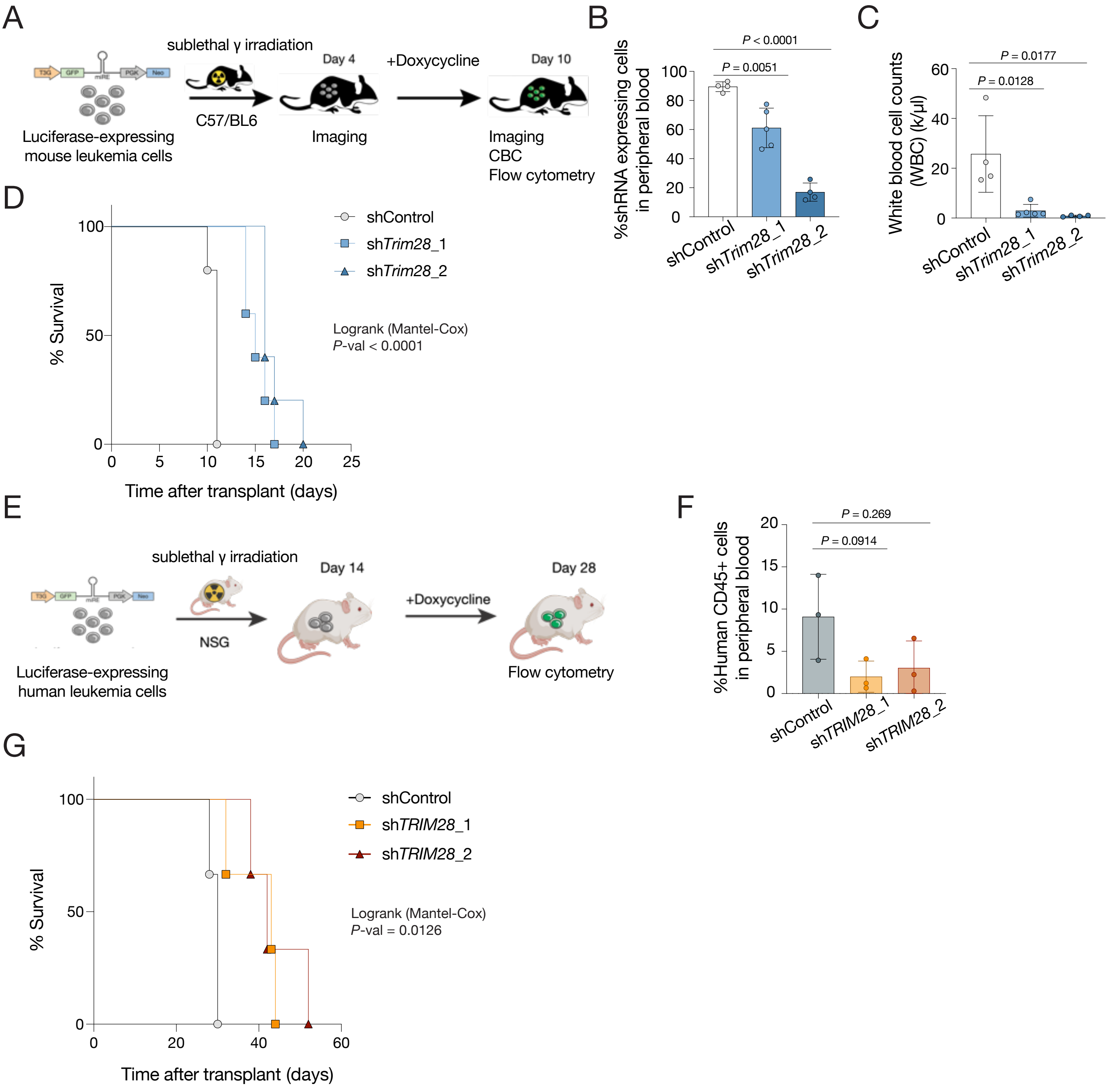


**Supplementary Figure 4 – *TRIM28* knockdown impairs leukemia progression *in vivo*. (A)** Schematic of the transplant of mouse MLL-AF9 leukemia cells expressing *Trim28* shRNAs. **(B)** Quantification of the proportion of GFP+ (shRNA-expressing) cells in peripheral blood at the terminal timepoint (n = 5). **(C)** Total white blood cell (WBC) counts at the terminal timepoint (n = 5). **(D)** Survival curves of syngeneic transplants of mouse MLL-AF9 leukemia cells engineered with doxycycline-inducible shRNAs. *P*-values were calculated using a log-rank test. **(E)** Schematic of the transplant of human MLL-AF9 leukemia cells (MOLM13) expressing *TRIM28* shRNAs. **(F)** Quantification of the proportion of CD45+ human leukemia cells in peripheral blood at the terminal timepoint (n = 3). **(G)** Survival curves of NSG mice transplanted with human MLL-AF9 leukemia cells (MOLM13) engineered with doxycycline-inducible shRNAs. *P*-value calculated by log-rank test.

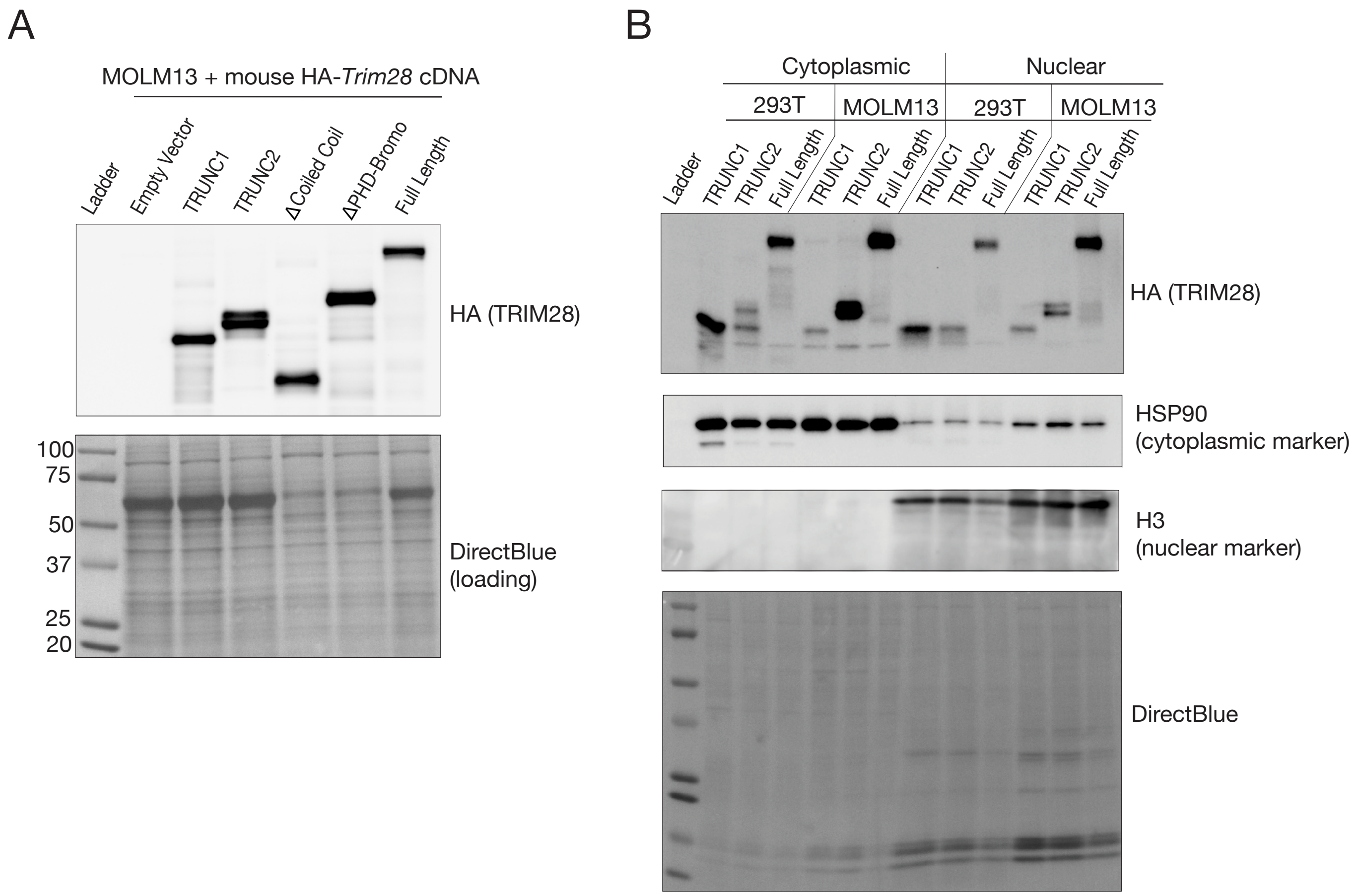


**Supplementary Figure 5 – Human leukemia cells constitutively express mouse TRIM28 protein. (A)** Immunoblot of human leukemia cells (MOLM13) constitutively expressing HA-*Trim28* cDNA constructs. The immunoblot for HA detects TRIM28 in each cell line; Direct Blue stain is shown below as a loading control. **(B)** Cellular fractionation followed by immunoblotting of HEK293T and MOLM13 cells expressing HA-*Trim28* cDNA constructs. Immunoblots show TRIM28 (top), HSP90 as a cytoplasmic marker (middle), and histone H3 as a nuclear marker (bottom); Direct Blue stain is shown below as a loading control.

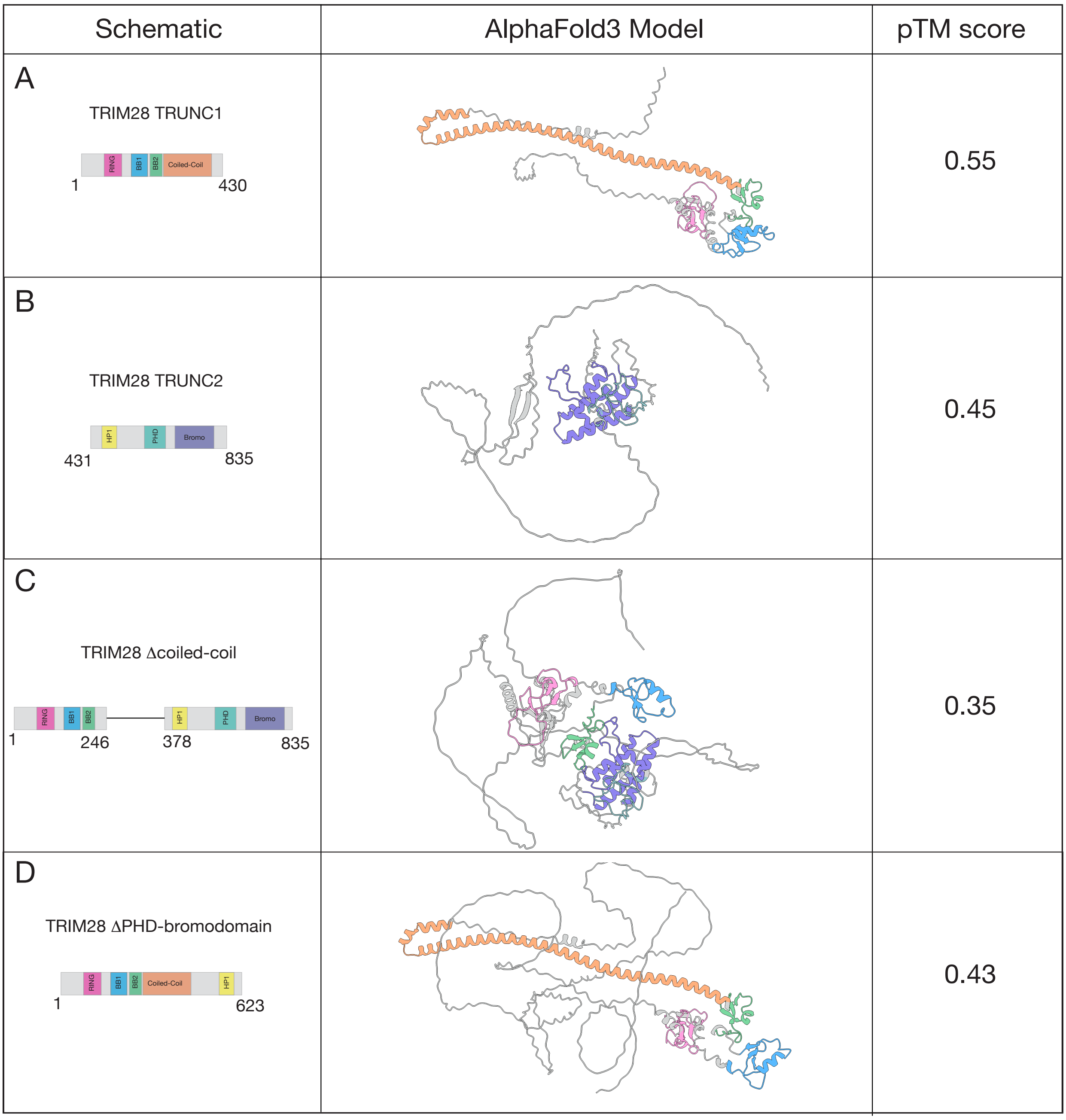


**Supplementary Figure 6 – TRIM28 coiled-coil domain is predicted to be required for structural integrity. (A)** (Left) Domain schematic of human TRIM28-TRUNC1, with amino acid positions shown below. (Middle) AlphaFold 3 model of human TRIM28-TRUNC1. (Right) Predicted template modeling score (pTM) of the AlphaFold 3 prediction. **(B)** (Left) Domain schematic of human TRIM28-TRUNC2, with amino acid positions shown below. (Middle) AlphaFold 3 model of human TRIM28-TRUNC2. (Right) Predicted template modeling score (pTM) of the AlphaFold 3 prediction. **(C)** (Left) Domain schematic of human TRIM28 coiled-coil internal deletion (Δcoiled-coil), with amino acid positions shown below. (Middle) AlphaFold 3 model of human TRIM28 Δcoiled-coil. (Right) Predicted template modeling score (pTM) of the AlphaFold 3 prediction. **(D)** (Left) Domain schematic of human TRIM28 PHD-bromodomain deletion (ΔPHD-bromodomain), with amino acid positions shown below. (Middle) AlphaFold 3 model of human TRIM28 ΔPHD-bromodomain. (Right) Predicted template modeling score (pTM) of the AlphaFold 3 prediction.

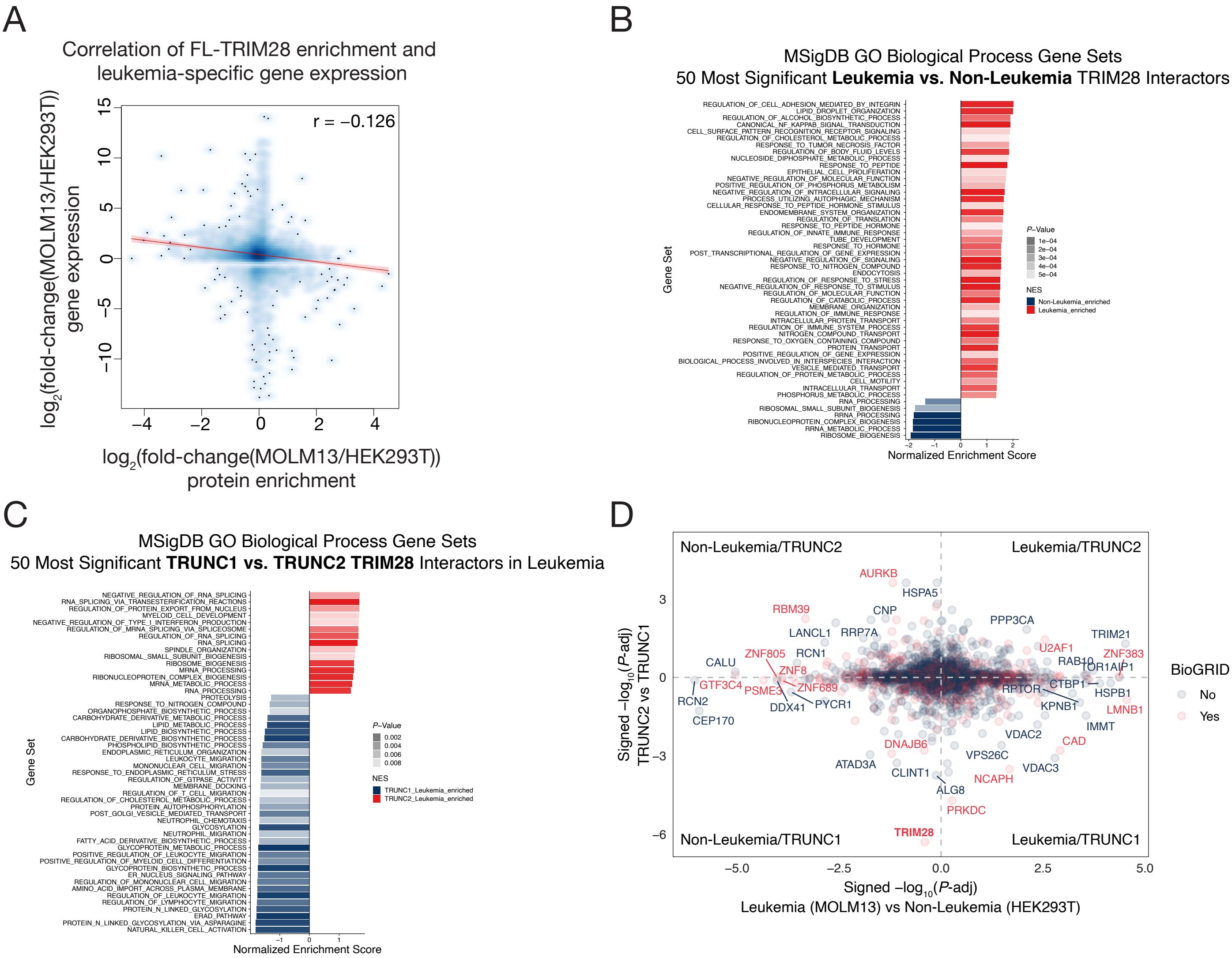


**Supplementary Figure 7 – Biological processes associated with the TRIM28 interactome in leukemia. (A)** Scatter plot with density overlay of $log_2$(fold-change) enrichment of full-length TRIM28 in human leukemia cells (MOLM13) compared to non-leukemia cells (HEK293T), determined by co-immunoprecipitation/mass spectrometry (co-IP/MS) (n = 4 replicates), versus $log_2$(fold-change) in gene expression determined by RNA-Seq in MOLM13 compared to HEK293T cells. Pearson correlation ($r$ = -0.126) is shown as a dark red line, with the 95% confidence interval shown as light red shading. **(B)** Gene Ontology (GO) analysis displaying the 50 most significantly enriched biological process gene sets (MSigDB GO BP) in the TRIM28 interactome in leukemia (MOLM13) versus non-leukemia (HEK293T) cells. Red bars represent pathways enriched in leukemia TRIM28 IP, whereas blue bars represent pathways enriched in non-leukemia TRIM28 IP. The $x$-axis shows the normalized enrichment score (NES); positive values indicate enrichment in leukemia, and negative values indicate enrichment in non-leukemia. Color saturation within each category (red or blue) reflects the $P$-value, with darker colors corresponding to more significant enrichment. **(C)** Gene Ontology (GO) analysis displaying the 50 most significantly enriched biological process gene sets (MSigDB GO BP) in the TRIM28-TRUNC2 interactome versus the TRIM28-TRUNC1 interactome in leukemia (MOLM13). Red bars represent pathways enriched in TRIM28-TRUNC2 IP, whereas blue bars represent pathways enriched in TRIM28-TRUNC1 IP. The $x$-axis shows the NES; positive values indicate enrichment in TRIM28-TRUNC2, and negative values indicate enrichment in TRIM28-TRUNC1. Color saturation within each category reflects the $P$-value, with darker colors corresponding to more significant enrichment. **(D)** Four-way comparison of protein enrichment in full-length TRIM28 immunoprecipitation in MOLM13 versus HEK293T ($x$-axis) and protein enrichment in TRIM28-TRUNC1 versus TRIM28-TRUNC2 ($y$-axis). Signed -$log_{10}$($P$-value) represents the measure of enrichment. Proteins annotated as TRIM28 interactors in the BioGRID database are shown in red, and proteins not annotated in BioGRID are shown in blue.

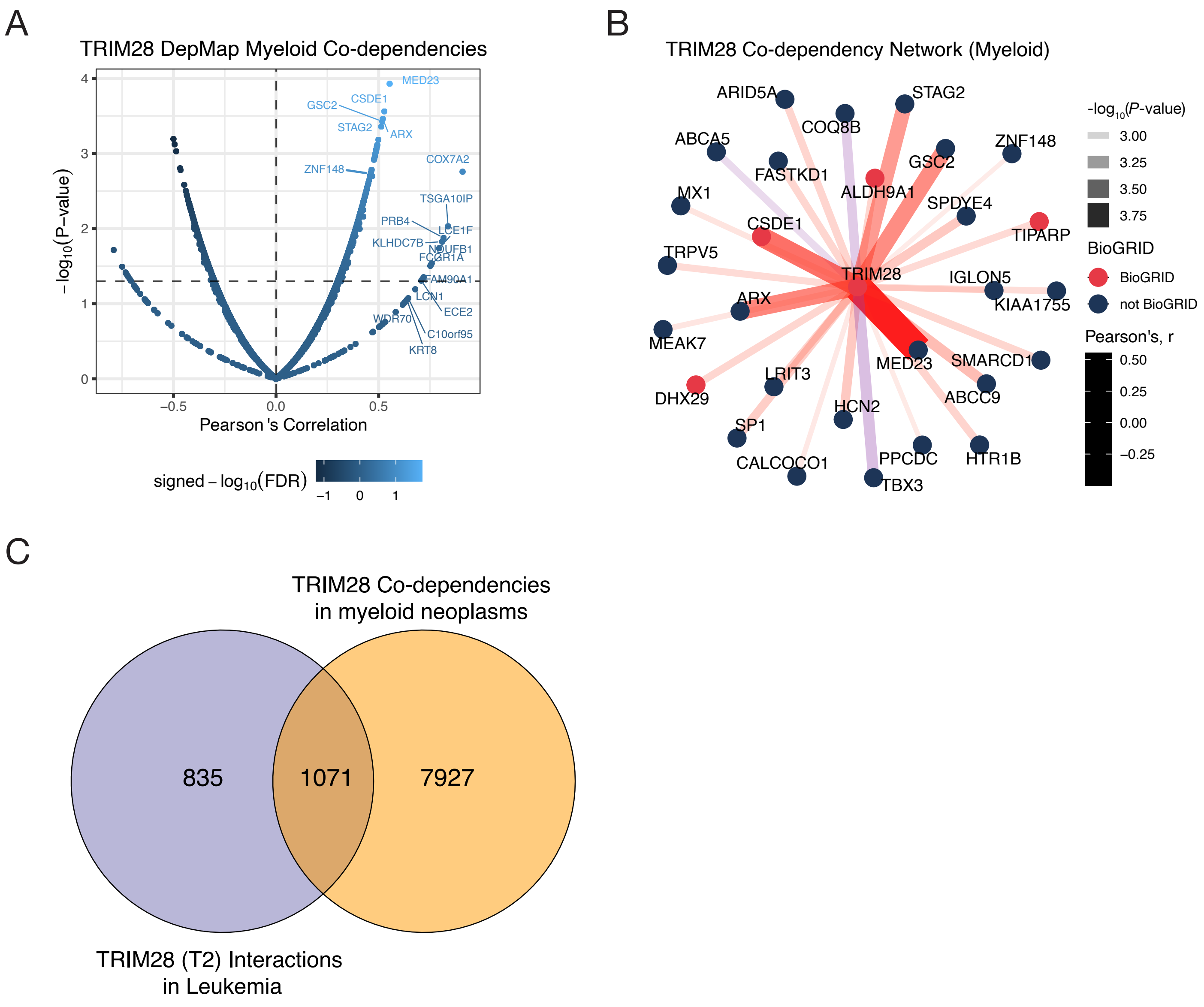


**Supplementary Figure 8 – Co-dependencies associated with TRIM28 in leukemia. (A)** Volcano plot of TRIM28 myeloid co-dependencies from DepMap. The *x*-axis shows the Pearson correlation coefficient for each co-dependent protein, and the *y*-axis shows -$\log_{10}$(*P*-value). Highly significant co-dependent proteins are in the top-right region of the plot. **(B)** TRIM28 myeloid co-dependencies shown as a network plot. BioGRID-annotated interactions are shown as blue nodes, and non-BioGRID interactions are shown as red nodes. Edge shading corresponds to the Pearson correlation coefficient, and color saturation corresponds to -$\log_{10}$(*P*-value). **(C)** Venn diagram of the co-dependencies shown in panels A and B, and the interaction partners of TRIM28 (TRUNC2) in leukemia.

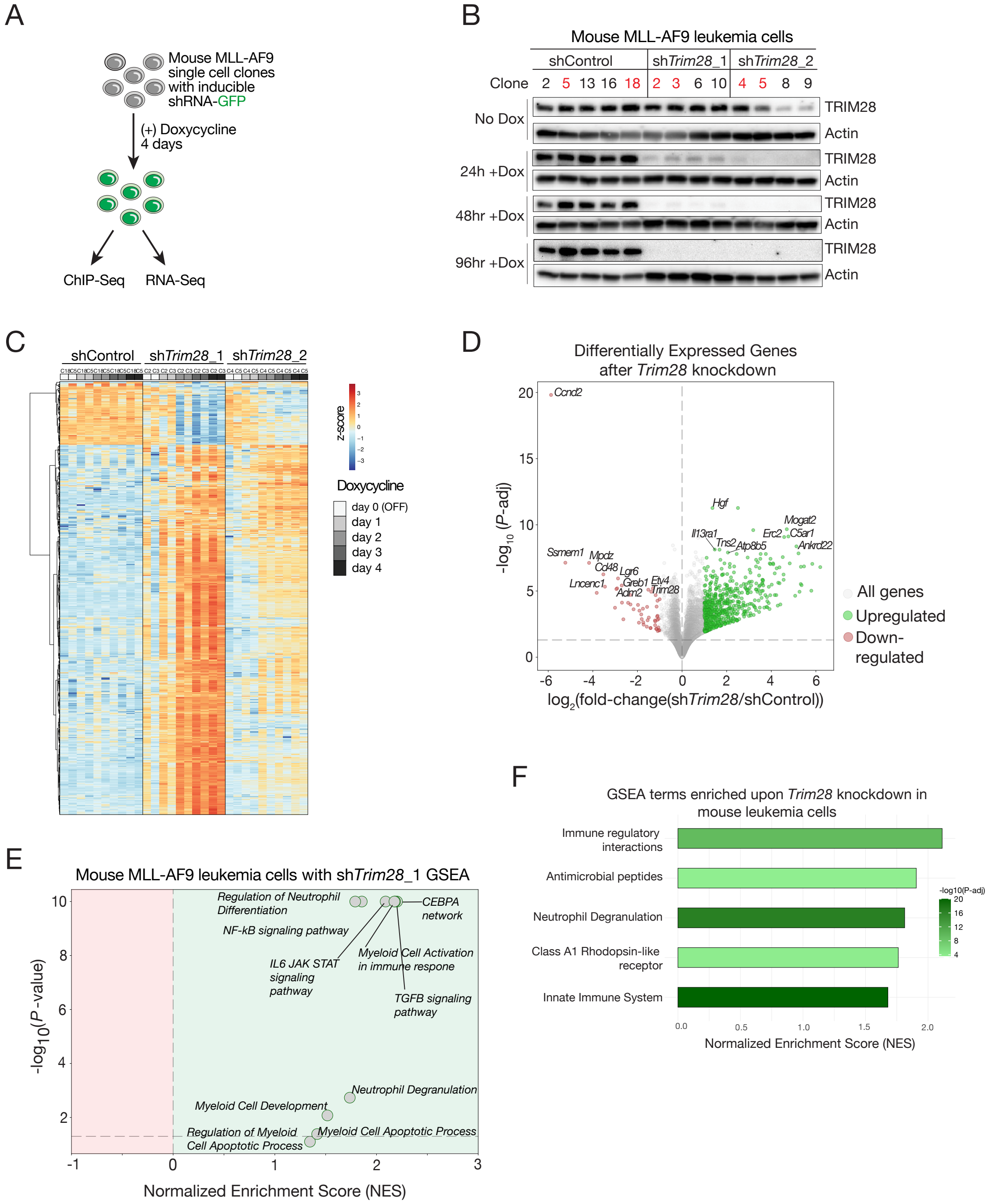


**Supplementary Figure 9 – Transcriptional profiling of *Trim28* loss in a mouse model of AML. (A)** Schematic of transcriptional and chromatin profiling in mouse MLL-AF9 leukemia cells after *Trim28* knockdown. **(B)** Immunoblot of mouse MLL-AF9 leukemia cells engineered with doxycycline-inducible shRNAs (non-targeting shControl; *Trim28*-targeting sh*Trim28*_1 and sh*Trim28*_2) co-expressing *GFP*. Cell pellets were collected before and 1, 2, and 4 days after shRNA induction. Four to five single-cell clones are shown for each shRNA condition, with clones selected for RNA-Seq highlighted in red. **(C)** Heatmap of differentially expressed genes in mouse leukemia cells with doxycycline-inducible shRNAs (sh*Trim28*_1, sh*Trim28*_2, or non-targeting shControl) before and 1, 2, 3, and 4 days after shRNA induction (two clones per shRNA condition). **(D)** Volcano plot of differentially expressed genes measured by RNA-Seq in mouse leukemia cells after 4 days of *Trim28* knockdown (effects of sh*Trim28*_1 and sh*Trim28*_2 averaged) compared to a non-targeting control (n = 3 replicates). Genes significantly upregulated following *Trim28* knockdown ($\log_2$(fold-change) > 1 and $-\log_{10}$(*P*-value) > 1.3) are shown in green, and genes significantly downregulated ($\log_2$(fold-change) < -1 and $-\log_{10}$(*P*-value) > 1.3) are shown in red. **(E)** Scatter plot of GSEA results for myeloid differentiation and apoptosis gene signatures among differentially expressed genes after 4 days of *Trim28* knockdown with sh*Trim28*_1. Gene ranks were calculated across replicates (n = 3). The *x*-axis shows the NES for each term, and the *y*-axis shows $-\log_{10}$(*P*-value); the most significantly enriched terms are in the top-right quadrant. **(F)** GSEA using Reactome gene sets after 4 days of *Trim28* knockdown in mouse leukemia cells (effects of sh*Trim28*_1 and sh*Trim28*_2 averaged). The top five most significantly enriched gene signatures are shown. The *x*-axis represents the normalized enrichment score (NES) for each term, and the shading of each bar represents $-\log_{10}$(*P*-value), with darker shading indicating more significant enrichment.

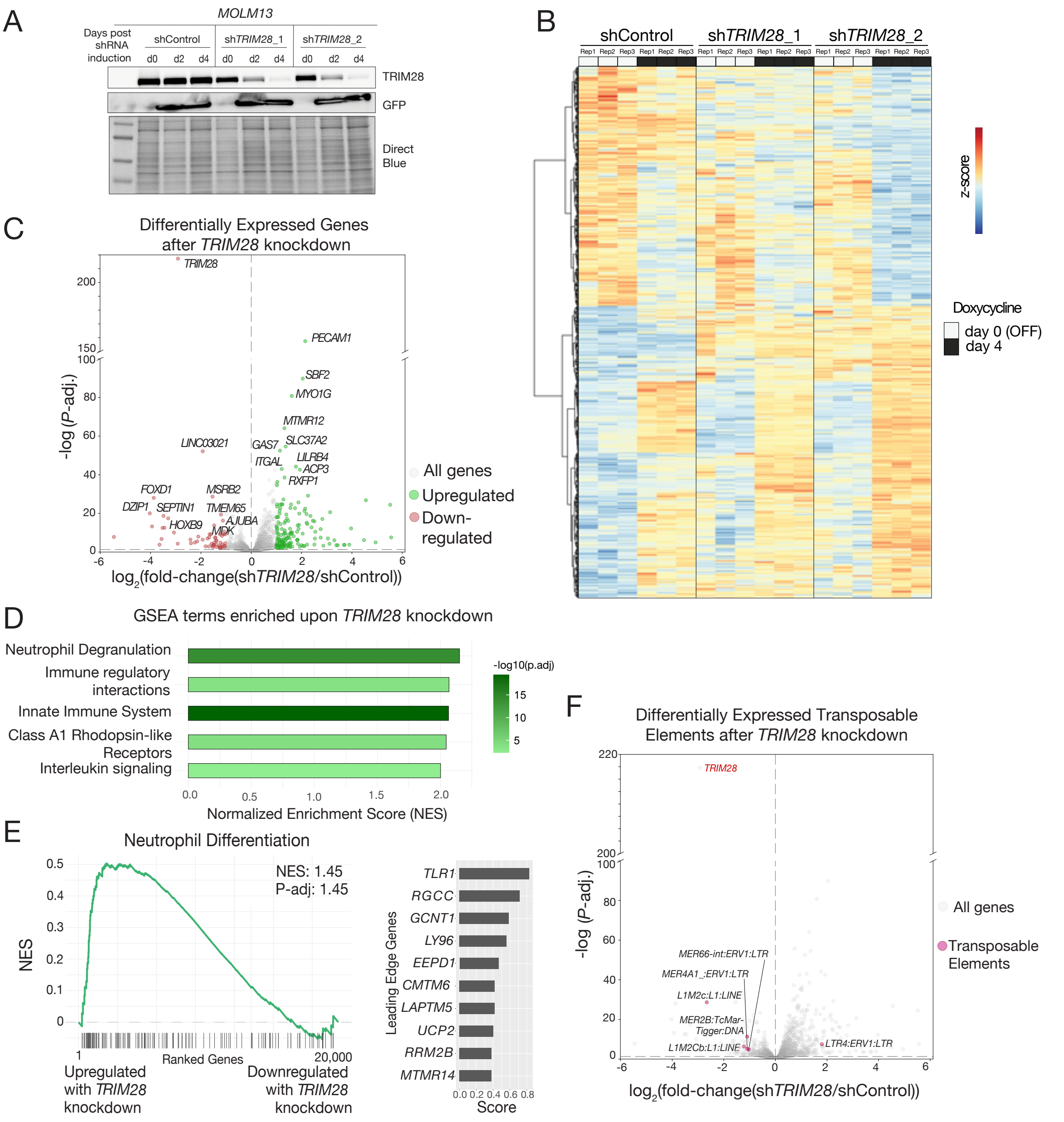


**Supplementary Figure 10 – Transcriptional profiling of *TRIM28* loss in a human model of AML. (A)** Immunoblot of human MLL-AF9 leukemia cells (MOLM13) engineered with doxycycline-inducible shRNAs (non-targeting shControl; *TRIM28*-targeting sh*TRIM28*_1 and sh*TRIM28*_2) co-expressing *GFP*. Cell pellets were collected before and 2 and 4 days after shRNA induction. TRIM28 (top) and GFP (middle) are shown, with Direct Blue staining below as a loading control. **(B)** Heatmap of differentially expressed genes upon *TRIM28* knockdown with doxycycline-inducible shRNAs targeting *TRIM28* or a non-targeting control before and 4 days after shRNA induction (n = 3 replicates). **(C)** Volcano plot of differentially expressed genes measured by RNA-Seq in MOLM13 cells after 4 days of *TRIM28* knockdown (sh*TRIM28*_1 or sh*TRIM28*_2) compared to a non-targeting control (n = 3 replicates), with transcriptional effects from both shRNAs averaged. Genes significantly upregulated following *TRIM28* knockdown ($log_2$(fold-change) > 1 and -$log_{10}$(*P*-value) > 1.3) are shown in green, and genes significantly downregulated ($log_2$(fold-change) < -1 and -$log_{10}$(*P*-value) > 1.3) are shown in red; the 10 most significantly upregulated and downregulated genes are annotated. **(D)** Gene set enrichment analysis (GSEA) using Reactome gene sets after 4 days of *TRIM28* knockdown in MOLM13 cells (effects of sh*TRIM28*_1 and sh*TRIM28*_2 averaged). The top five most significantly enriched gene signatures are shown. The *x*-axis shows the normalized enrichment score (NES) for each term, and the shading of each bar represents -$log_{10}$(*P*-value), with darker shading indicating more significant enrichment. **(E)** GSEA plot showing enrichment of a neutrophil differentiation gene signature (GSE27786) following 4 days of *TRIM28* knockdown (effects of sh*TRIM28*_1 and sh*TRIM28*_2 averaged) in MOLM13 cells. The NES and *P*-value are indicated, and the top 10 leading-edge genes are shown in the bar plot to the right. **(F)** Volcano plot of differentially expressed transposable elements measured by RNA-Seq in MOLM13 cells after 4 days of *TRIM28* knockdown (sh*TRIM28*_1 or sh*TRIM28*_2) compared to a non-targeting control (n = 3 replicates), with effects from both shRNAs averaged. Transposable elements significantly altered (up- or downregulated) following *TRIM28* knockdown (|$log_2$(fold-change)| > 1 and -$log_{10}$(*P*-value) > 1.3) are shown in pink and annotated.

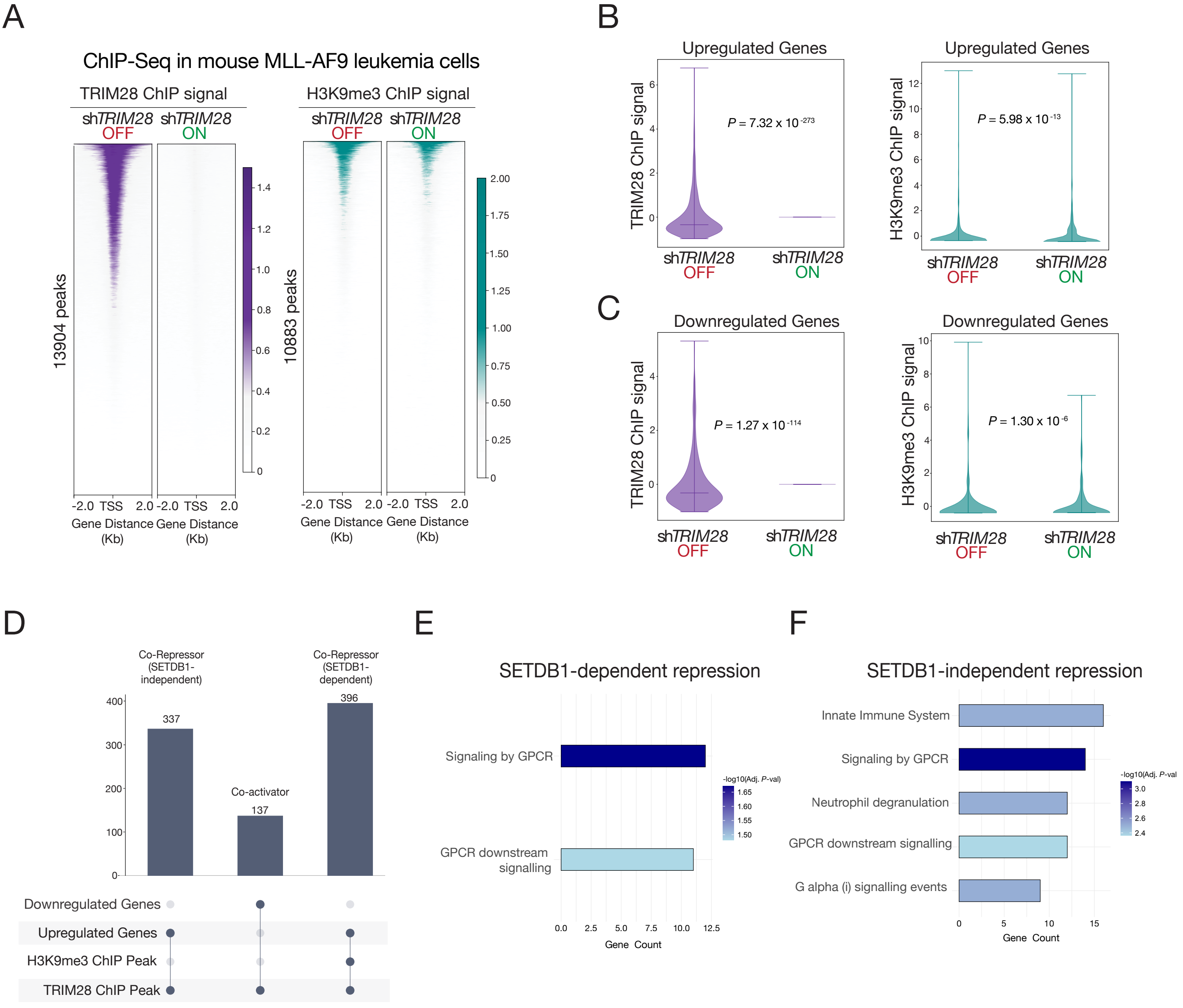


**Supplementary Figure 11 – TRIM28 regulates neutrophil genes through a SETDB1-independent mechanism.** **(A)** (Left) Genome-wide profiling of TRIM28 chromatin occupancy by ChIP-Seq in mouse MLL-AF9 leukemia cells before and after *Trim28* knockdown (sh*Trim28*_1), with peaks sorted by TRIM28 signal. (Right) Genome-wide profiling of H3K9me3 chromatin occupancy by ChIP-Seq in the same cells before and after *Trim28* knockdown, with peaks co-sorted by TRIM28 signal. **(B)** Violin plots of TRIM28 ChIP signal (left) and H3K9me3 ChIP signal (right) at genes upregulated upon *Trim28* knockdown, defined by RNA-Seq (n = 1,716), before and 4 days after sh*Trim28*_1 induction. **(C)** Violin plots of TRIM28 ChIP signal (left) and H3K9me3 ChIP signal (right) at genes downregulated upon *Trim28* knockdown, defined by RNA-Seq (n = 754), before and 4 days after sh*Trim28*_1 induction. **(D)** Upset plot showing overlap among genes with TRIM28 ChIP peaks, genes with H3K9me3 ChIP peaks, TRIM28-upregulated genes, and TRIM28-downregulated genes. **(E)** Reactome pathway analysis of the top 100 TRIM28-upregulated genes that have both TRIM28 and H3K9me3 ChIP peaks, representing SETDB1-dependent repression. The *x*-axis shows the number of genes in each pathway, and the shading of each bar represents -$\log_{10}$(*P*-value), with darker shading indicating more significant enrichment. **(F)** Reactome pathway analysis of the top 100 TRIM28-upregulated genes with TRIM28 ChIP peaks but no H3K9me3 ChIP peaks, representing SETDB1-independent repression. The *x*-axis shows the number of genes in each pathway, and the shading of each bar represents -$\log_{10}$(*P*-value), with darker shading indicating more significant enrichment.

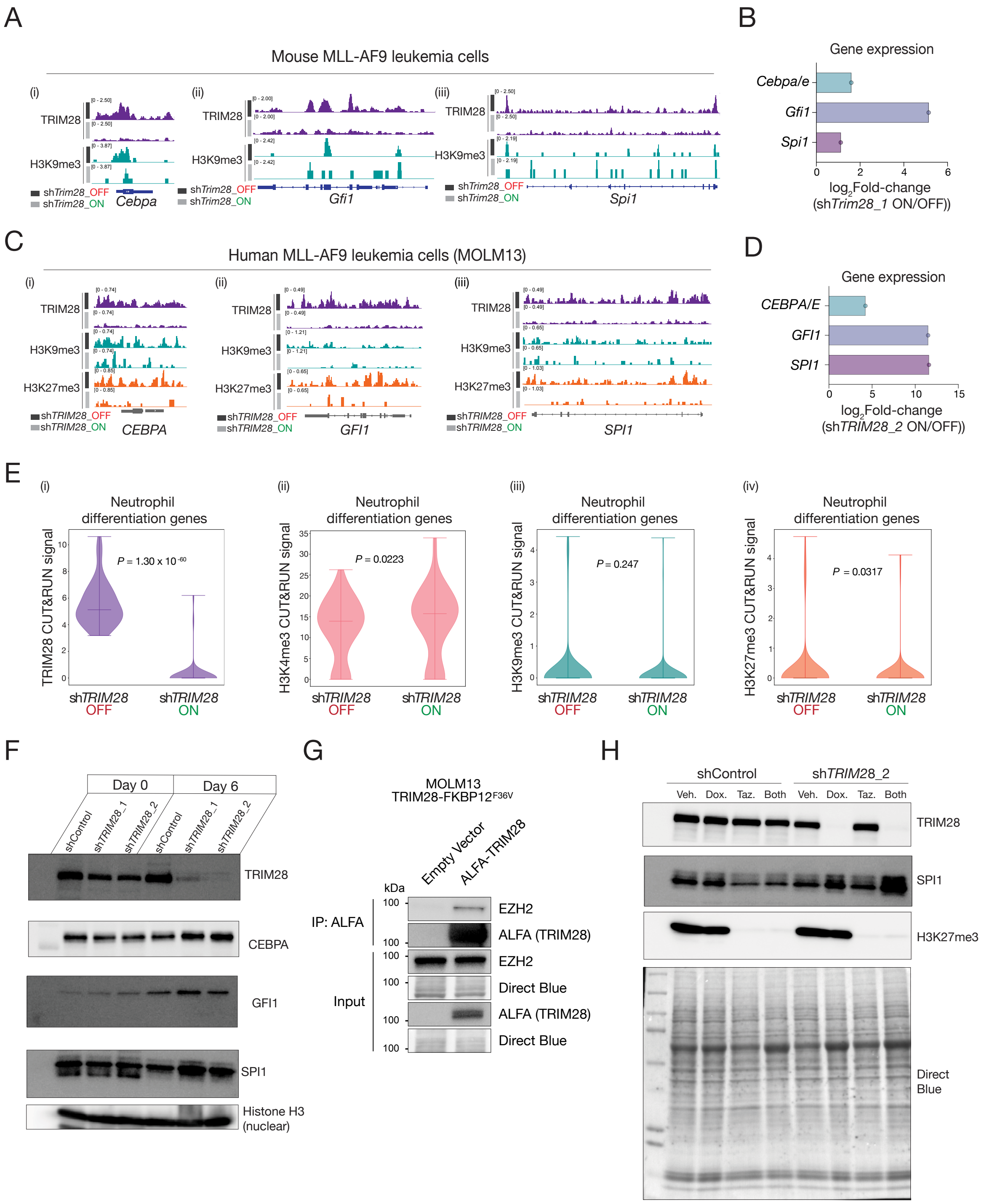


**Supplementary Figure 12 – TRIM28 localizes to the promoters of neutrophil differentiation genes. (A)** TRIM28 (purple) and H3K9me3 (teal) ChIP-Seq signal at the (i) *Cebpa*, (ii) *Gfi1*, and (iii) *Spi1* promoters before and after *Trim28* knockdown (sh*Trim28*_1). **(B)** Gene expression of *Cebpa*, *Gfi1*, and *Spi1*, measured by RNA-Seq in mouse leukemia cells before and after *Trim28* knockdown (sh*Trim28*_1). **(C)** TRIM28 (purple), H3K9me3 (teal), and H3K27me3 (orange) CUT&RUN signal at the (i) *CEBPA*, (ii) *GFI1*, and (iii) *SPI1* promoters before and after *TRIM28* knockdown (sh*TRIM28*_2). **(D)** Gene expression of *CEBPA*, *GFI1*, and *SPI1*, measured by RNA-Seq in human leukemia cells before and after *TRIM28* knockdown (sh*TRIM28*_2). **(E)** Violin plots of (i) TRIM28, (ii) H3K4me3, (iii) H3K9me3, and (iv) H3K27me3 CUT&RUN signal at neutrophil differentiation genes (n = 88), before and 4 days after sh*TRIM28*_2 induction. **(F)** Nuclear extraction followed by immunoblotting in human leukemia cells before and 6 days after *TRIM28* knockdown (sh*TRIM28*_1 or sh*TRIM28*_2). The immunoblot shows TRIM28 (top), CEBPA, GFI1, SPI1 (middle panels), and H3 as a nuclear marker (bottom). **(G)** (Top) Immunoprecipitation for ALFA tag followed by immunoblot showing EZH2 and ALFA tag in human leukemia cells expressing ALFA-tagged full-length TRIM28 or empty vector. (Bottom) Input lysate immunoblot showing EZH2 and ALFA tag. Direct Blue staining is shown below each immunoblot as a loading control. **(H)** Nuclear extraction followed by immunoblotting in human leukemia cells treated for 6 days with vehicle, *TRIM28* knockdown by shRNA (sh*TRIM28*_2), EZH2 inhibitor tazemetostat (Taz), or the combination of sh*TRIM28*_2 and Taz. The immunoblot shows TRIM28 (top), SPI1 (middle), and H3K27me3 (bottom), with Direct Blue staining below as a loading control.

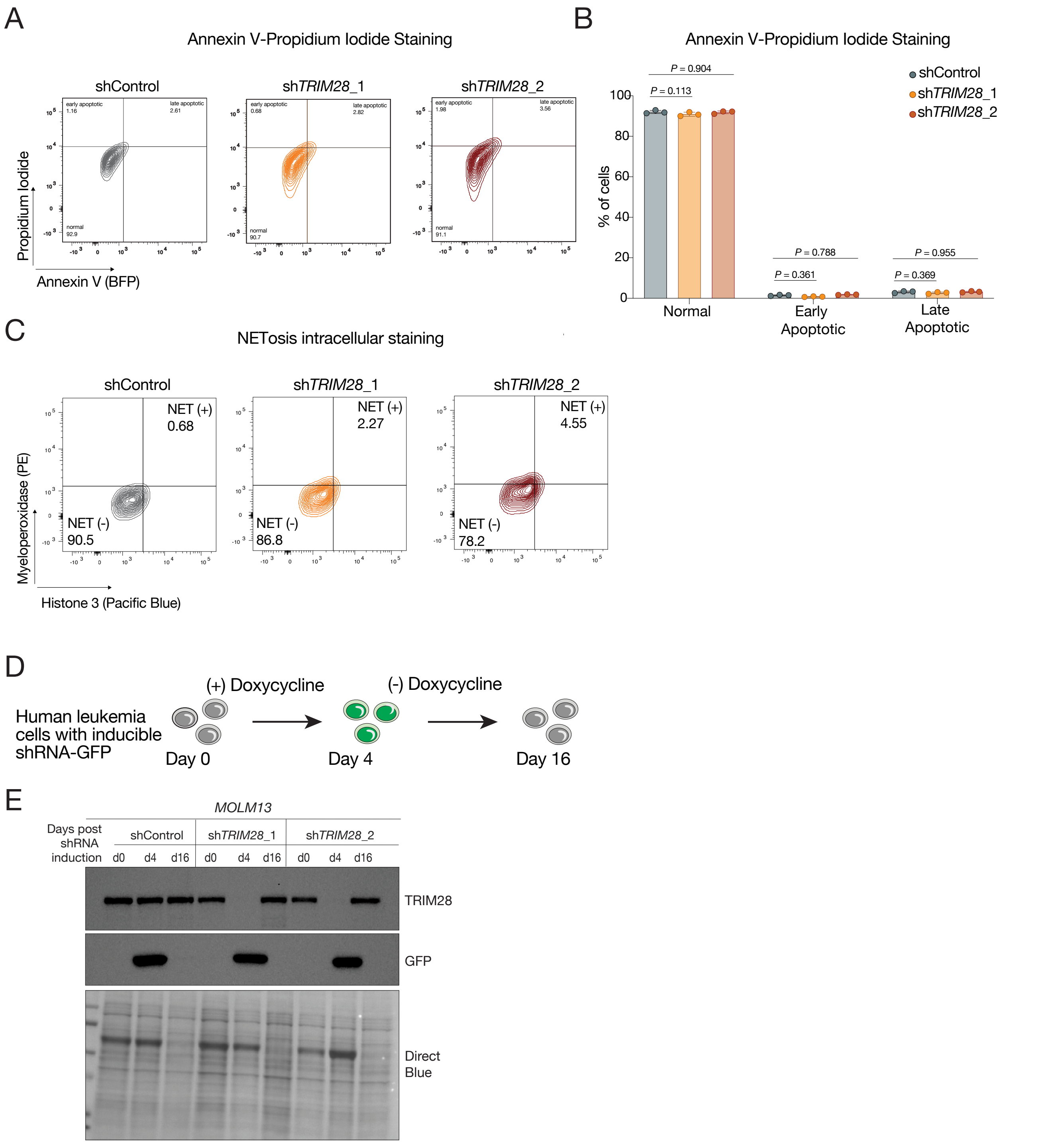


**Supplementary Figure 13 – *TRIM28* loss does not lead to cell death via apoptosis. (A)** Representative plot of Annexin V-propidium iodide staining to assess apoptosis in human leukemia cells (MOLM13) after 12 days of doxycycline-induced shRNA expression (non-targeting shControl; *TRIM28*-targeting sh*TRIM28*_1 and sh*TRIM28*_2). **(B)** Quantification of Annexin V-propidium iodide staining in MOLM13 cells after 12 days of doxycycline-induced shRNA expression (n = 3 replicates). **(C)** Representative cell surface MPO and histone H3 staining to assess NETosis in MOLM13 cells after 12 days of doxycycline-induced shRNA expression. **(D)** Schematic of the reversibility experiment in which MOLM13 cells engineered with doxycycline-inducible shRNAs were treated with doxycycline for 4 days, maintained on doxycycline until a neutrophil differentiation phenotype was established (day 12), and then cultured without doxycycline for 4 additional days. Cell pellets were collected before shRNA induction, during shRNA expression, and after shRNA withdrawal. **(E)** Immunoblot of MOLM13 cells before shRNA induction, 4 days after doxycycline treatment, and 4 days after doxycycline withdrawal. Immunoblots show TRIM28 (top), GFP (middle), and Direct Blue staining below as a loading control.

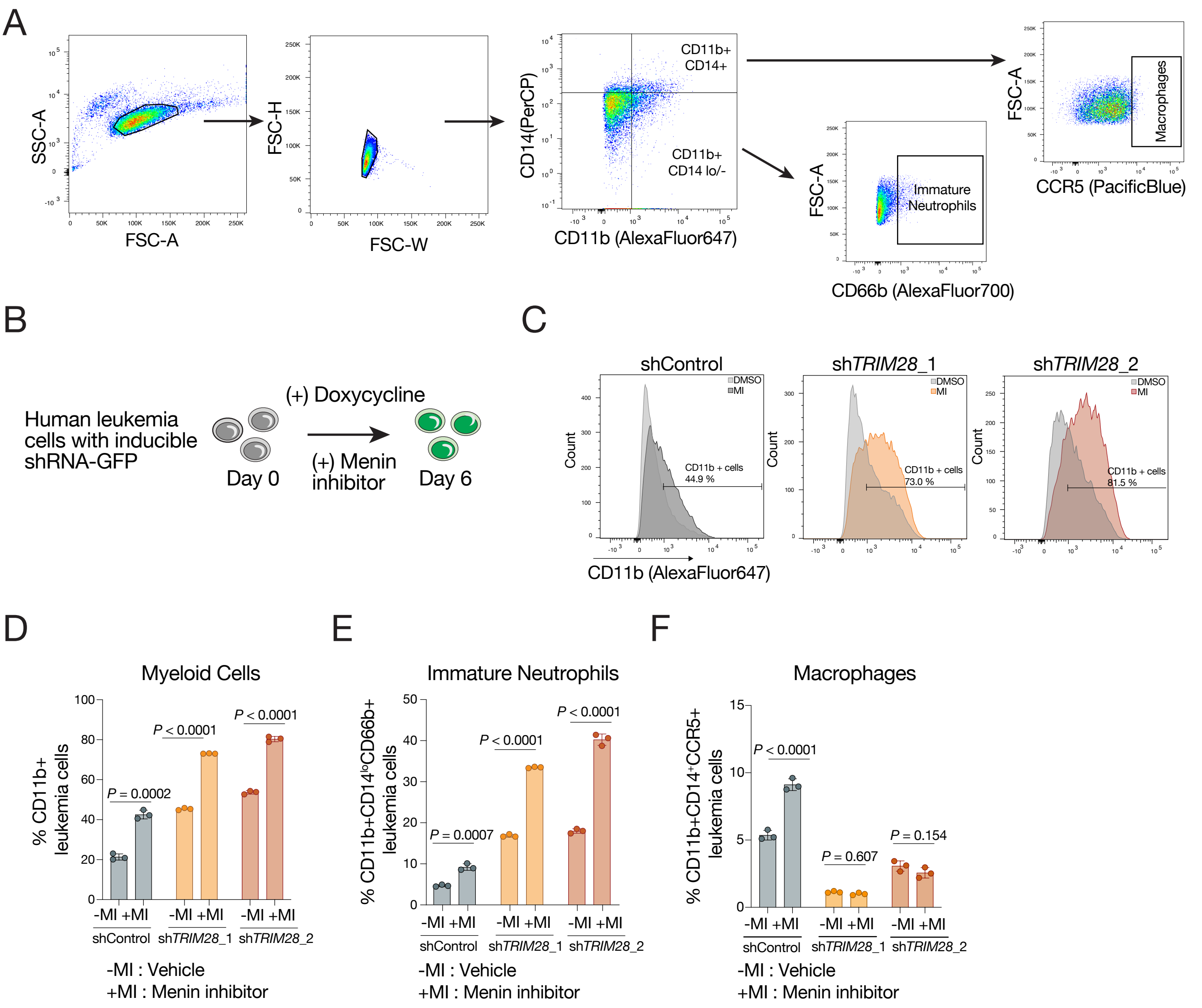


**Supplementary Figure 14 – Menin-MLL inhibitor amplifies phenotypic consequences of *TRIM28* loss. (A)** Gating schematic for cell-surface staining followed by flow cytometry to distinguish neutrophil and macrophage populations. **(B)** Schematic of the dual *TRIM28* knockdown and Menin inhibitor treatment experiment. **(C)** Representative flow cytometry plot of CD11b$^+$ human leukemia cells after simultaneous *TRIM28* knockdown and Menin inhibitor treatment ($IC_{50}$, 6 days), gated relative to vehicle control. **(D)** Proportion of myeloid (CD11b$^+$) cells (gated relative to vehicle control) under the same conditions as in Supplementary Figure 14C (n = 3 replicates). **(E)** Proportion of immature neutrophils (CD11b$^+$, CD14$^{lo}$, CD66b$^+$) (gated relative to vehicle control) under the same conditions as in Supplementary Figure 14C (n = 3 replicates). **(F)** Proportion of macrophages (CD11b$^+$, CD14$^+$, CCR5$^+$) under the same conditions as in Supplementary Figure 14C (n = 3 replicates).

A

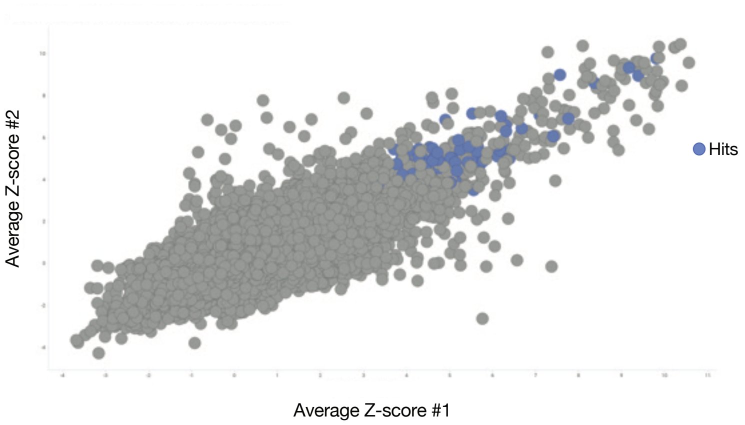


**Supplementary Figure 15 – High-throughput small-molecule screening for TRIM28 PHD-bromodomain binders. (A)** Scatter plot showing the average Z-scores across technical replicates for each compound in the original screen (*x*-axis) and the repeat screen (*y*-axis). Hit compounds are highlighted in blue; hits were defined as having *Z*-scores > 3 and signal-to-background ratios > 1.3 across all four replicates and being absent from control small-molecule microarray screens lacking the TRIM28 PHD-bromodomain.

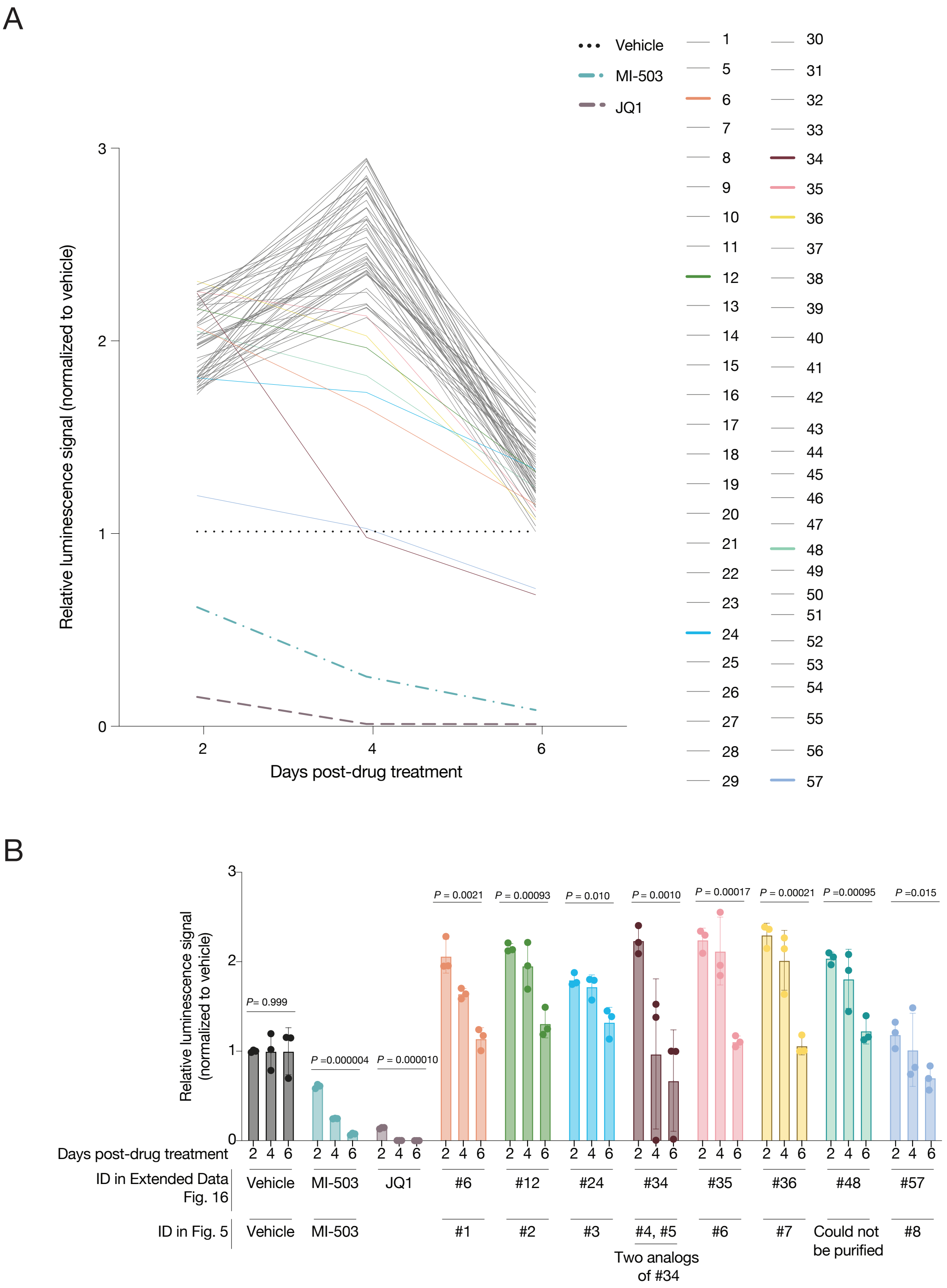


**Supplementary Figure 16 – TRIM28 PHD-bromodomain binding molecules inhibit proliferation of human leukemia cells. (A)** Proliferation curves for MOLM13 human leukemia cells treated with each small-molecule inhibitor; cell viability measured every 2 days for 6 days, normalized to vehicle control (DMSO), and mean viability for each compound was plotted (n = 3 replicates). Menin inhibitor (MI-503) and BRD4 inhibitor (JQ1) were included as positive controls. **(B)** (Top) Proliferation of MOLM13 cells treated with each TRIM28 PHD-bromodomain hit compound, with viability measured every 2 days for 6 days, normalized to vehicle control (DMSO) and displayed as bar graphs (n = 3 replicates). Menin inhibitor (MI-503) and BRD4 inhibitor (JQ1) were included as positive controls. (Bottom) Table listing IDs for each small-molecule hit in the initial screen shown in Supplementary Figure 16A and in the secondary screen shown in Figure 5B-C.

A

| KI-T28 | Chembridge ID | Structure |
|---|---|---|
| 01 | 82561814 | 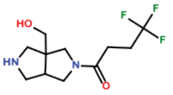 |
| 02 | 48799456 | 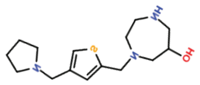 |
| 03 | 7977502 | 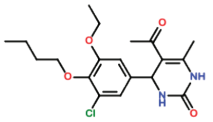 |
| 04 | 16797437 | 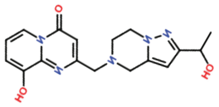 |
| 05 | 95333716 | 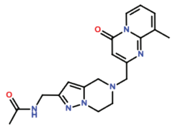 |
| 06 | 94536620 | 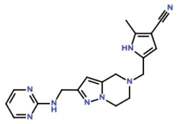 |
| 07 | 98007575 | 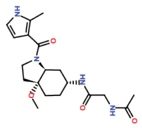 |
| 08 | 31295389 | 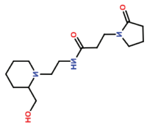 |

B

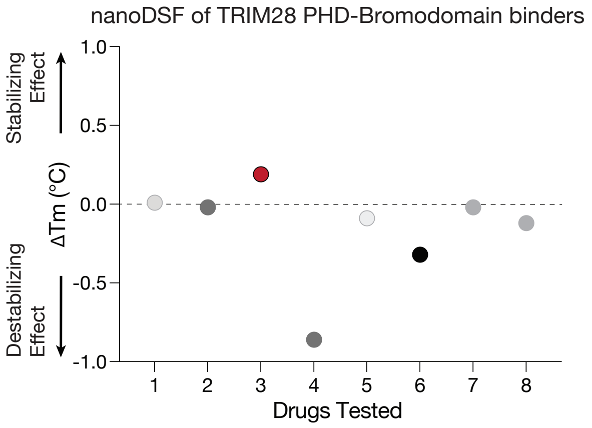


**Supplementary Figure 17 – TRIM28 PHD-bromodomain binders alter protein stability. (A)** Table listing the KI-T28 inhibitor IDs used in Figure 5, the corresponding ChemBridge catalog IDs, and the chemical structure of each compound. **(B)** Change in melting temperature (ΔTm), measured by nanoDSF, for each TRIM28 small-molecule candidate. A positive ΔTm indicates stabilization of TRIM28 upon small-molecule binding, whereas a negative ΔTm indicates destabilization.

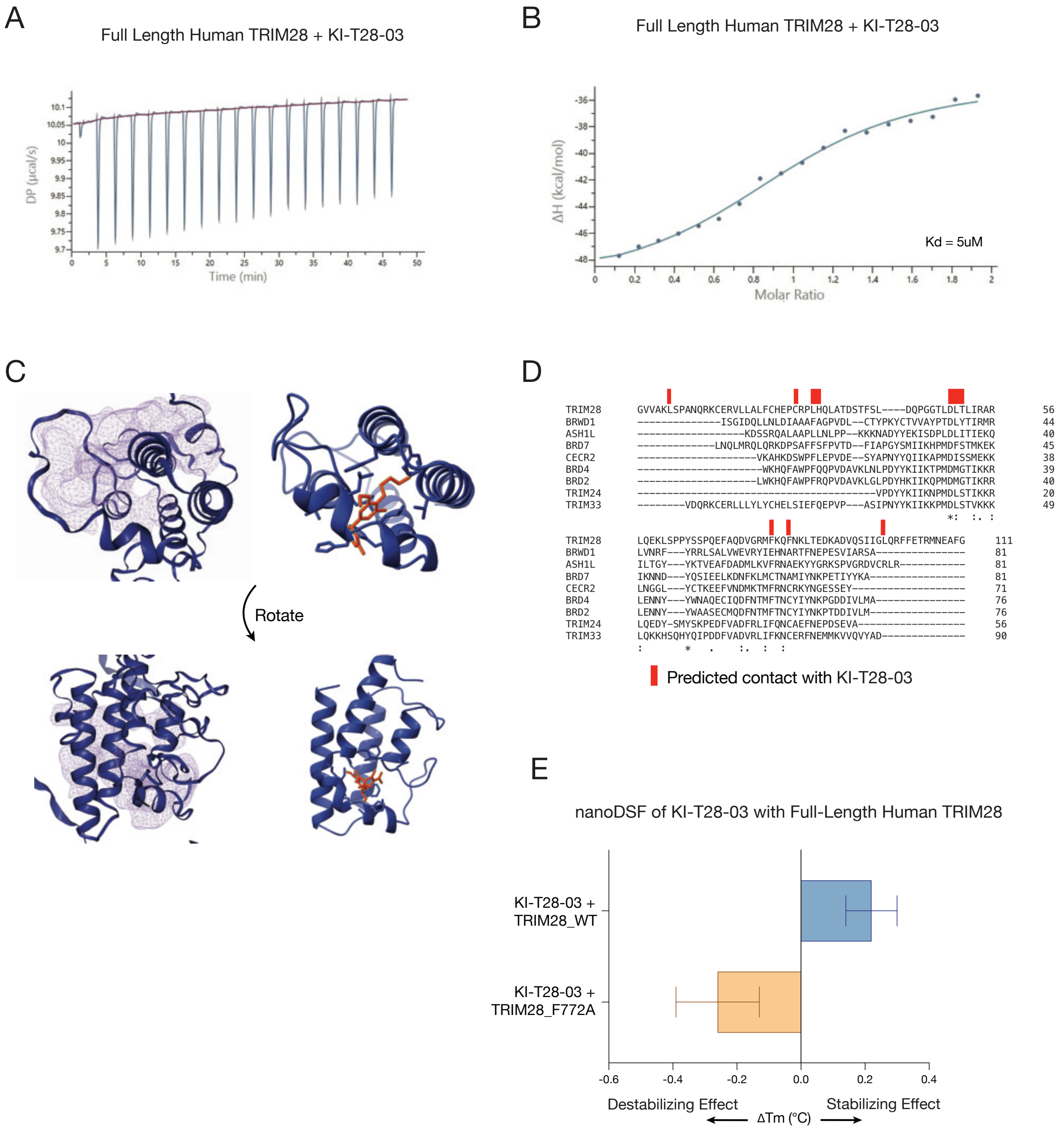


**Supplementary Figure 18 – KI-T28-03 binds the TRIM28 bromodomain with high affinity. (A)** Isothermal titration calorimetry (ITC) differential power (DP) time-course traces showing raw injection heats for a titration of KI-T28-03 into full-length TRIM28, after baseline correction. **(B)** Integrated heat curve plotting binding enthalpy as a function of molar ratio. The solid line represents a nonlinear least-squares fit to a one-site binding model used to calculate KD, yielding a fitted KD of 5 µM. **(C)** AlphaFold 3 model of full-length human TRIM28 highlighting the predicted KI-T28-03 binding pocket, shown in two orientations, with TRIM28 in blue and KI-T28-03 in orange. **(D)** Structure-based sequence alignment of TRIM28 and selected bromodomains, with residues predicted to contact KI-T28-03 marked in red above the alignment. **(E)** Change in melting temperature (ΔTm), measured by nanoDSF, for wild-type and F772A full-length TRIM28 upon binding to KI-T28-03. A positive ΔTm indicates a stabilizing effect on TRIM28 upon small-molecule binding, whereas a negative ΔTm indicates a destabilizing effect (n = 3 technical replicates).

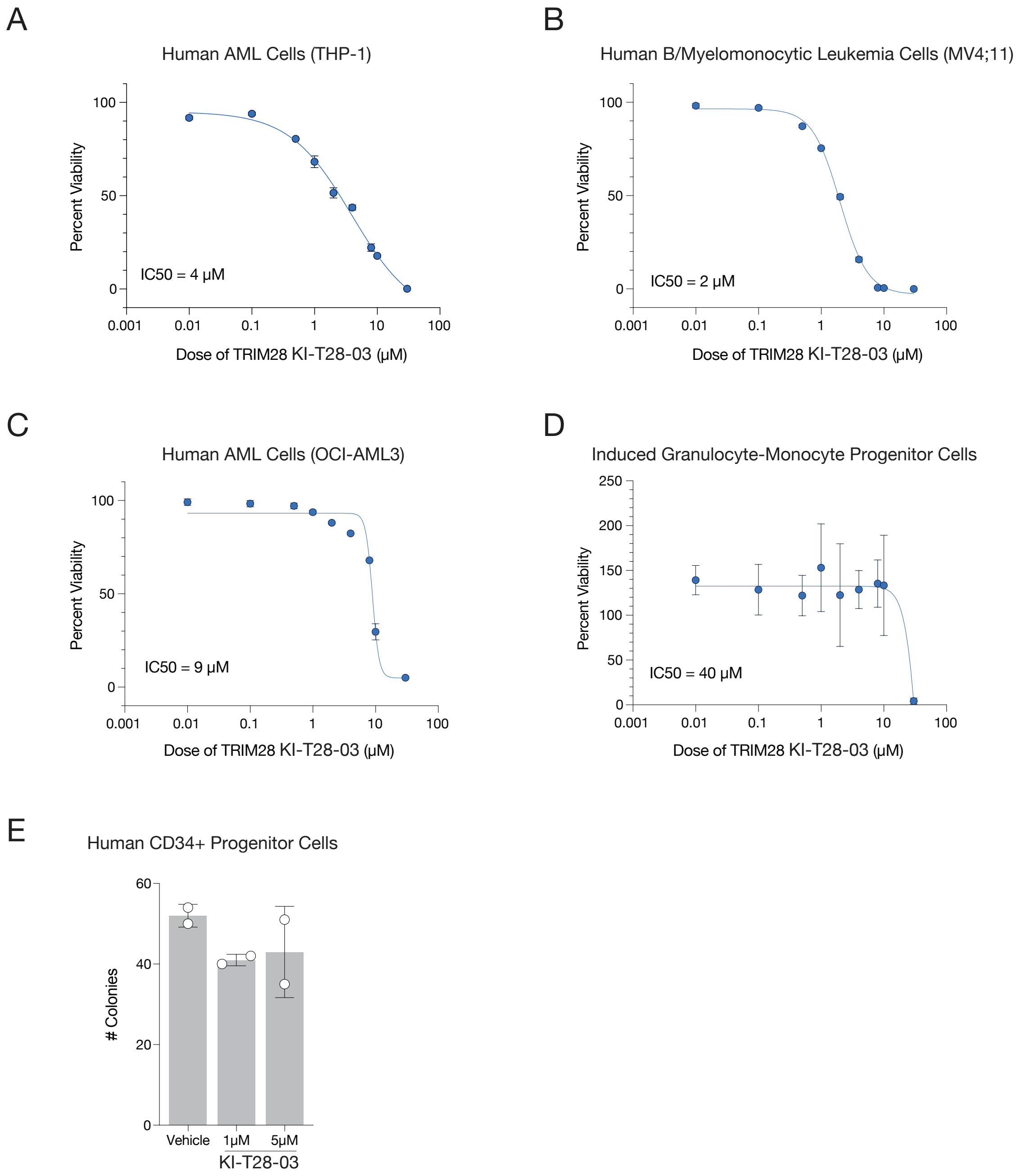


**Supplementary Figure 19 – TRIM28 inhibitor KI-T28-03 selectively impairs growth of multiple human leukemia cell lines. (A-C)** Viability of human leukemia cell lines across six concentrations of KI-T28-03. The cell line name and key driver mutation are indicated above each plot. Viability was measured, normalized to vehicle control (DMSO), and fitted with a nonlinear regression curve with $IC_{50}$ indicated in the lower left of each panel (n = 3 replicates). **(D)** Viability of induced granulocyte-monocyte progenitors (iGMPs) across six concentrations of KI-T28-03, normalized to vehicle control (DMSO), and fitted with a nonlinear regression curve with $IC_{50}$ indicated (n = 3 replicates). **(E)** Bar plot showing the number of colony-forming units generated from human $CD34^{+}$ progenitor cells treated with 1 μM or 5 μM KI-T28-03, with DMSO-treated cells serving as a negative control.

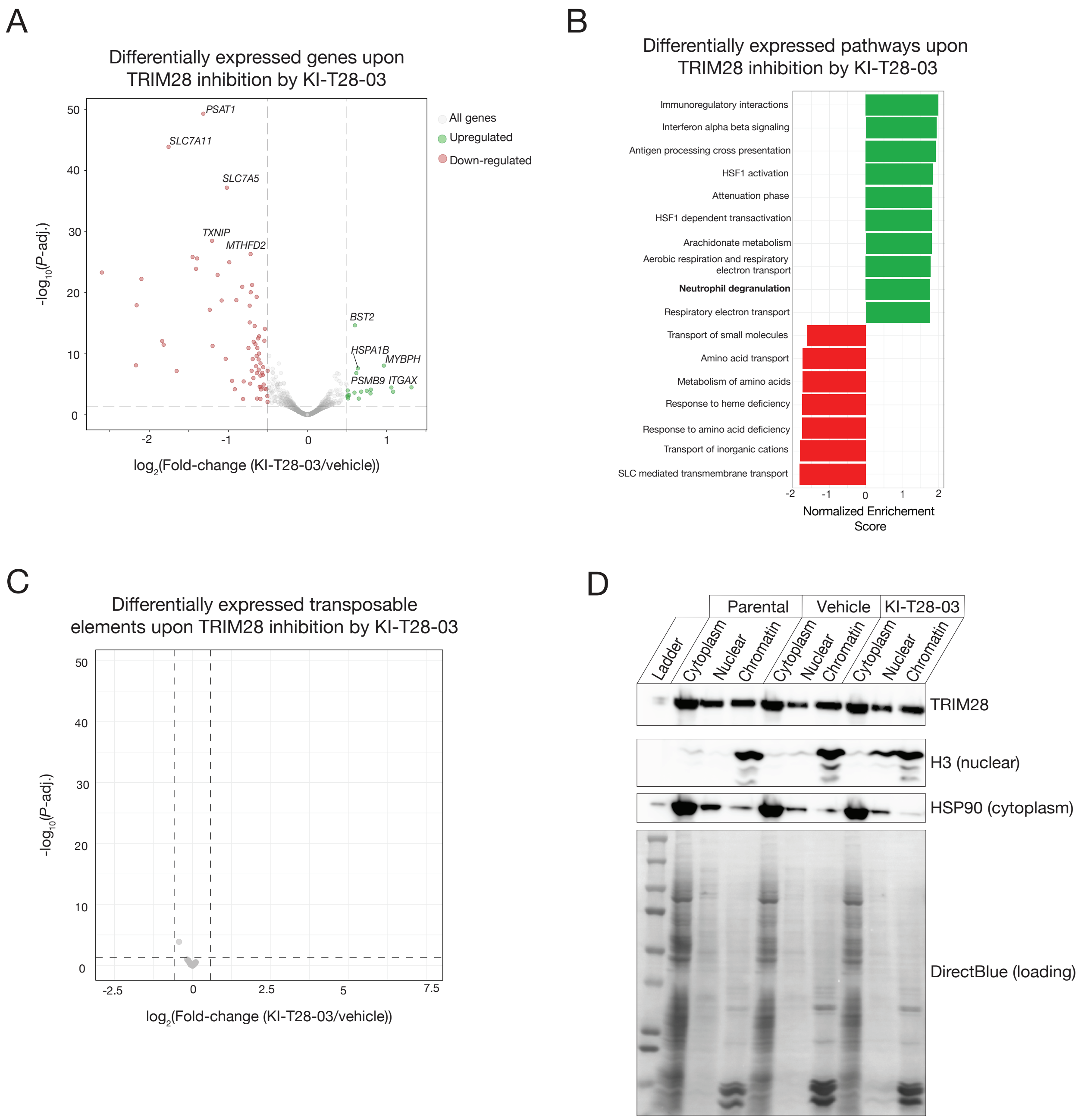


**Supplementary Figure 20 – TRIM28 inhibitor KI-T28-03 reprograms transcription and induces neutrophil differentiation. (A)** Volcano plot of differentially expressed genes measured by RNA-Seq in human leukemia cells after 4 days of TRIM28 small-molecule inhibition compared to vehicle control (n = 3 replicates). Genes significantly upregulated following KI-T28-03 treatment ($log_2$(fold-change) > 1 and $-log_{10}$(*P*-value) > 1.3) are shown in green, and genes significantly downregulated ($log_2$(fold-change) < -1 and $-log_{10}$(*P*-value) > 1.3) are shown in red. **(B)** GSEA plot showing the 20 most significantly enriched reactome gene sets (MSigDB Reactome) in human leukemia cells treated with KI-T28-03 compared vehicle control (DMSO). Green bars represent pathways upregulated in response to KI-T28-03 (normalized enrichment score > 1), and red bars represent pathways downregulated (normalized enrichment score < -1). **(C)** Volcano plot of differentially expressed transposable elements measured by RNA-Seq in human leukemia cells after 4 days of TRIM28 small-molecule inhibition compared to vehicle control (n = 3 replicates). **(D)** Biochemical cellular fractionation followed by immunoblotting of MOLM13 cells treated with vehicle control (DMSO) or 5 µM KI-T28-03. Immunoblots show TRIM28 (top), H3 as a nuclear control (middle), and HSP90 as a cytoplasmic control (bottom), with Direct Blue staining below as a loading control.

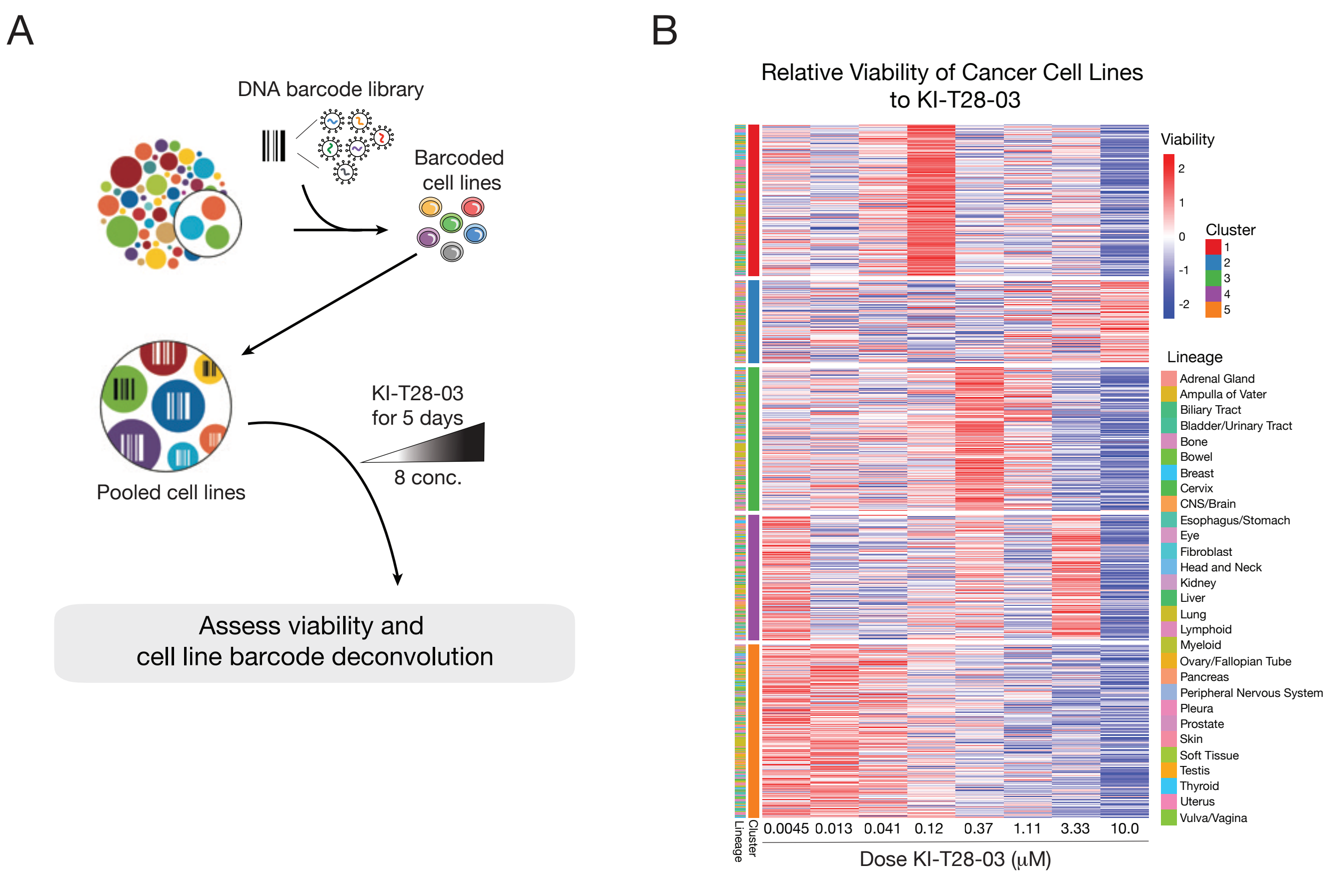


**Supplementary Figure 21 – PRISM pooled screening identifies cancer lineages sensitive to TRIM28 inhibitor KI-T28-03. (A)** Schematic of the Broad Institute PRISM workflow, in which barcoded cancer cell lines are pooled, treated with KI-T28-03 at eight concentrations for 5 days, and then assessed for viability by barcode deconvolution. **(B)** Heatmap showing relative viability of 917 cancer cell lines across increasing concentrations of KI-T28-03. Cell lines are grouped into five clusters by *K*-means clustering, and the lineage of each cell line is indicated by the color-coded annotation bar on the left.
